# Corticostriatal Maldevelopment in the R6/2 Mouse Model of Juvenile Huntington’s Disease

**DOI:** 10.1101/2024.10.15.618500

**Authors:** Carlos Cepeda, Sandra M. Holley, Joshua Barry, Katerina D. Oikonomou, Vannah-Wila Yazon, Allison Peng, Deneen Argueta, Michael S. Levine

## Abstract

There is a growing consensus that brain development in Huntington’s disease (HD) is abnormal, leading to the idea that HD is not only a neurodegenerative but also a neurodevelopmental disorder. Indeed, structural and functional abnormalities have been observed during brain development in both humans and animal models of HD. However, a concurrent study of cortical and striatal development in a genetic model of HD is still lacking. Here we report significant alterations of corticostriatal development in the R6/2 mouse model of juvenile HD. We examined wildtype (WT) and R6/2 mice at postnatal (P) days 7, 14, and 21. Morphological examination demonstrated early structural and cellular alterations reminiscent of malformations of cortical development, and *ex vivo* electrophysiological recordings of cortical pyramidal neurons (CPNs) demonstrated significant age- and genotype-dependent changes of intrinsic membrane and synaptic properties. In general, R6/2 CPNs had reduced cell membrane capacitance and increased input resistance (P7 and P14), along with reduced frequency of spontaneous excitatory and inhibitory synaptic events during early development (P7), suggesting delayed cortical maturation. This was confirmed by increased occurrence of GABA_A_ receptor-mediated giant depolarizing potentials at P7. At P14, the rheobase of CPNs was significantly reduced, along with increased excitability. Altered membrane and synaptic properties of R6/2 CPNs recovered progressively, and by P21 they were similar to WT CPNs. In striatal medium-sized spiny neurons (MSNs), a different picture emerged. Intrinsic membrane properties were relatively normal throughout development, except for a transient increase in membrane capacitance at P14. The first alterations in MSNs synaptic activity were observed at P14 and consisted of significant deficits in GABAergic inputs, however, these also were normalized by P21. In contrast, excitatory inputs began to decrease at this age. We conclude that the developing HD brain is capable of compensating for early developmental abnormalities and that cortical alterations precede and are a main contributor of striatal changes. Addressing cortical maldevelopment could help prevent or delay disease manifestations.

**Graphical Abstract:** 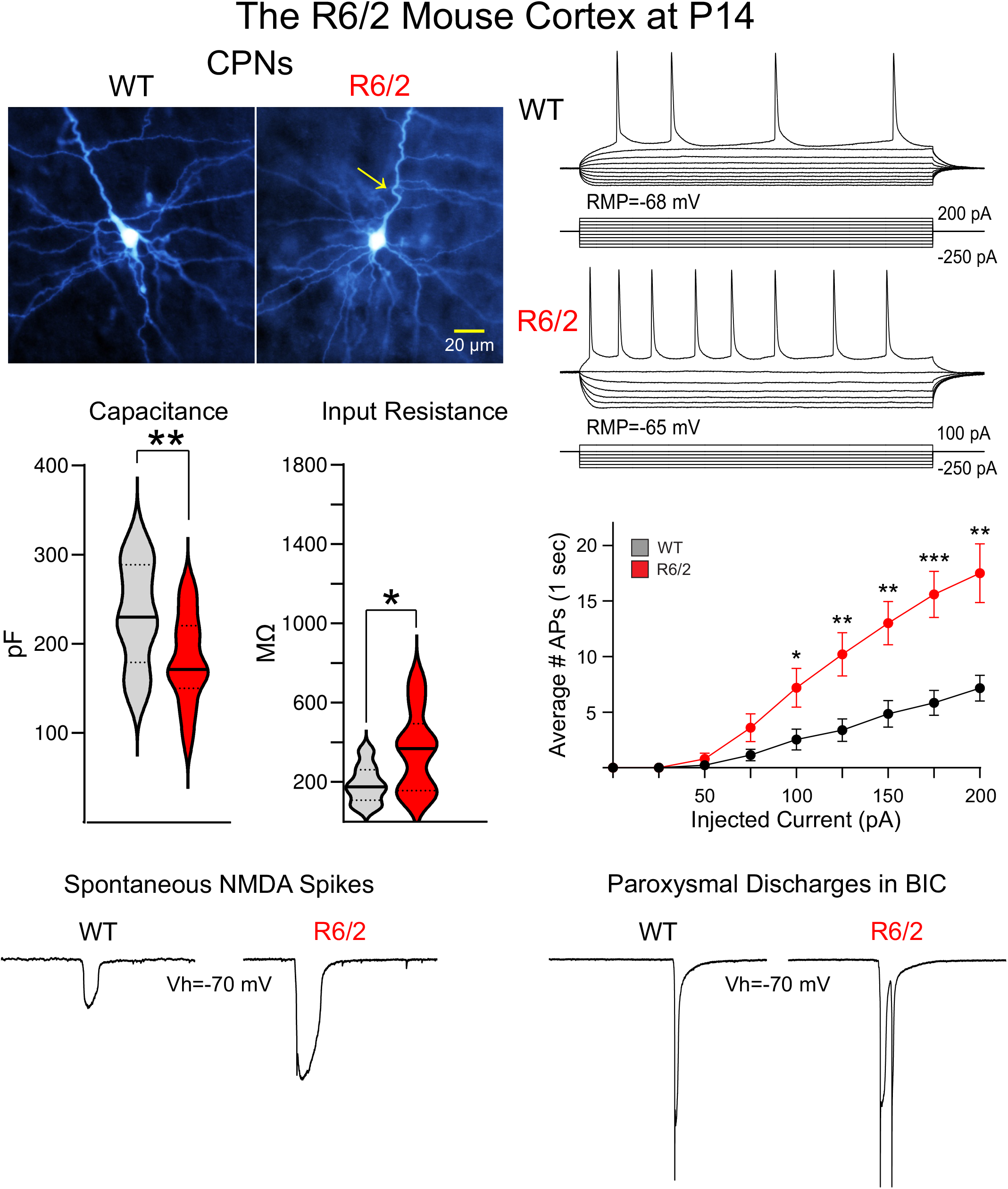

## Introduction

Huntington’s disease (HD) is typically an adult-onset neurodegenerative disorder generally starting after 30 years of age, and much later in patients with short CAG repeat expansions (>36 but <45) (MacDonald and Gusella, 1996). Severity of symptoms is dictated by the number of CAG repeats, as well as various genetic and environmental modifiers (Pengo and Squitieri, 2024). In general, the greater the number of CAG repeats the earlier the onset of HD symptoms (Andrew et al., 1993; Squitieri et al., 2006; Stine et al., 1993). When the number of CAG repeats exceeds 60, another form of HD emerges, known as juvenile HD (JHD or Westphal variant). Pediatric or childhood-onset HD also occurs when the expansion is >80 CAG repeats (Letort and Gonzalez-Alegre, 2013). JHD, with age of onset at <20 years, is extremely rare (less than 10% of HD cases), has a rapid progression, and preferentially associates with paternal transmission (Telenius et al., 1993). Individuals with pediatric or JHD display phenotypic manifestations that are at variance from those of adult-onset HD. In those cases, chorea rarely occurs and instead, rigidity similar to that found in Parkinson’s disease is observed (Jervis, 1963).

In JHD, particularly in pediatric HD cases, epileptic seizures are a common symptom, whereas in adult-onset HD seizures almost never occur. Besides the epileptic syndrome, children with HD display dystonia, mental retardation, hyperactivity, and aggressive behavior (Oosterloo et al., 2024). Some of these children also have microcephaly, suggesting a neurodevelopmental rather than or in addition to a neurodegenerative disorder (Gonzalez-Alegre and Afifi, 2006; Letort and Gonzalez-Alegre, 2013; Ratie and Humbert, 2024). Notably, in HD patients and in genetic animal models, structural changes in diverse brain regions are now well documented (Barnat et al., 2020; Blockx et al., 2012; Consortium, 2017; Molero et al., 2016; Nopoulos et al., 2011; Tereshchenko et al., 2019; Zhang et al., 2020). A neuropathologic study in a large cohort of HD cases reported a significant increase in developmental malformations, in particular periventricular nodular heterotopias, compared to control cases (Hickman et al., 2021), suggesting cortical migration deficits. In agreement, we previously demonstrated the existence of structural changes in the cortex of R6/2 mice, a *bona fide* model of JHD (Cepeda et al., 2019). Our findings revealed close similarities between structural cortical alterations seen in JHD and focal cortical dysplasia (FCD), a malformation of cortical development that leads to hyperexcitability and epileptic seizures (Blumcke et al., 2011; Cepeda et al., 2006). In JHD and in R6/2 mice epileptic seizures are a common phenotype. Although seizures do not seem to occur in adult-onset HD or in slowly progressing HD genetic models (e.g., zQ175, BACHD, or YAC128), they have been reported in an enhanced zQ175 mouse model (Southwell et al., 2016).

Huntingtin (HTT) is essential for normal cortical development and its absence becomes embryonically lethal (Duyao et al., 1995; Reiner et al., 2003; Zeitlin et al., 1995). Although normal levels of mutant (m)HTT are sufficient to rescue neurogenic function (White et al., 1997), both the absence of wildtype HTT or the presence of mHTT can affect neuronal migration and polarization (Barnat et al., 2017; Godin et al., 2010; Molina-Calavita et al., 2014). HTT is expressed very early in developing brains (Bhide et al., 1996) and in R6/2 and YAC128 models, mHTT aggregates have been observed in developing axonal tracts (Osmand et al., 2016). This suggests that synaptic transmission along the corticostriatal pathway could be affected very early during brain development.

In a recent report, abnormal synaptic communication was observed in the cerebral cortex of a knock-in mouse model of HD (Hdh^Q111/Q7^) during the first week of development (Braz et al., 2022). Early deficits in glutamatergic activity could be rescued by treatment with an ampakine.

However, changes in GABAergic synaptic activity and how glutamatergic synaptic abnormalities affect striatal neuron development were not investigated. This is important as it has been shown that both cortical and striatal development are altered in JHD (Tereshchenko et al., 2019) and in mouse models (Lebouc et al., 2020). In addition, there is little information on cortical and striatal development in the most commonly used model of HD, the R6/2 mouse line. In an effort to determine which region shows the first morphological and electrophysiological alterations during early brain development, here we examined cortical and striatal development of R6/2 mice in tandem. Our findings demonstrate significant alterations in both cortex and striatum, albeit cortical abnormalities are more severe and appear to precede those in striatum. Some of the material in the present study was published in preliminary form (Cepeda et al., 2019).

## Methods

Experimental procedures were performed in accordance with the United States Public Health Service Guide for Care and Use of Laboratory Animals and were approved by the Institutional Animal Care and Use Committee at the University of California, Los Angeles (UCLA). Mice were obtained from our breeding colony and every effort was made to minimize pain, discomfort, and the number of mice used. The number of mice used for ex vivo studies are shown in Suppl. Tables I-III. Animal housing conditions were maintained under a standard 12/12 h light/dark cycle (light cycle starting at 6 A.M. and ending at 6 P.M.) and at a temperature of 20–26°C. The animals had *ad libitum* access to food and water. All experiments were performed using the R6/2 mouse line (CAG repeat length: 158 ± 1.8) and wildtype (WT) littermates. For these mice, WT male C57BL/6xCBA mice were crossed with WT female C57BL/6xCBA mice that had transplanted R6/2 ovaries (B6CBA-Tg(HDexon1)62Gpb/3J, RRID: IMSR_JAX:006494, The Jackson Laboratory). Most experiments used mice at three postnatal (P) ages (in days), P7±1, P14±2, and P21±3, except for a few additional morphological studies that used mice at P30±3 and P60±4 (for the sake of simplicity, groups are referred to as P7, P14, P21, P30 and P60). No effort was made to separate the mice by gender as it is difficult to determine sex at P7 and P14, thus data from male and female animals, including the P21 and older groups, were pooled. In order to limit the number of mice used, whenever possible, multiple experiments were performed using corticostriatal brain slices from the same mouse.

### Cortical Morphology

R6/2 mice and WT littermates at P14, P21, P30, and P60 were perfused and the brains stained for the specific neuronal marker NeuN using standard procedures. In brief, brains were fixed with phosphate-buffered 4% paraformaldehyde, cryoprotected in 20% phosphate-buffered sucrose, cut with a cryostat at 30 μm and mounted on coated slides. For NeuN staining, the free-floating sections were processed following the Mouse on Mouse Immunodetection Kit protocol (Vector Laboratories) and incubated in NeuN antisera (mouse anti-neuronal nuclei, Chemicon International, Cat. no. MAB377) at a 1:1000 dilution. Sections were further processed with a 3,3’-diaminobenzidine tetrahydrochloride (DAB) substrate kit (Vector ABC Elite Mouse kit) and developed in 0.5 mg/ml DAB and 0.01% hydrogen peroxide. The tissue was then mounted on coated slides, dehydrated and cover slipped with DePeX mounting medium.

### Slice electrophysiology

Detailed procedures have been published previously (Andre et al., 2011; Cepeda et al., 2008; Cepeda et al., 2013). In brief, mice were deeply anesthetized with isoflurane and sacrificed. The brains were dissected and immediately placed in oxygenated ice-cold high sucrose slice solution containing the following (in mM): 208 sucrose, 2.5 KCl, 1.25 NaH_2_PO_4_, 26 NaHCO_3_, 1.3 MgCl_2_, 8 MgSO_4_, and 10 glucose. Coronal slices containing both cerebral cortex and striatum were used to examine changes in membrane properties and spontaneous synaptic activity of CPNs from the motor cortex (M1, mostly layers II-III) and MSNs from dorsolateral striatum in HD mice and WT littermates. Slices were cut (300 µm) and transferred to an incubating chamber containing artificial cerebrospinal fluid (ACSF) (in mM): 130 NaCl, 3 KCl, 1.25 NaH_2_PO_4_, 26 NaHCO_3_, 2 CaCl_2_, 2 MgCl_2_, and 10 glucose, oxygenated with 95% O_2_-5% CO_2_, pH 7.2–7.4, 290–310 mOsm, 25±2°C) at 32°C for 30 min. Slices continued to recover at room temperature for an additional 30 min before recordings. All slices were visualized with infrared differential interference contrast (IR-DIC) microscopy and cells were selected for voltage or current clamp recordings. Most experiments were performed in voltage clamp mode. For these recordings, the patch pipette (3-5 MΩ impedance) contained the following solution (in mM): 125 Cs-methanesulfonate, 4 NaCl, 3 KCl, 1 MgCl_2_, 5 MgATP, 9 EGTA, 8 HEPES, 0.5 GTP, 10 phosphocreatine, 0.1 leupeptin (pH 7.25-7.3, osmolality, 280-290 mOsm). In all experiments, only cells with access resistances <25 MΩ were recorded. Cells were initially voltage clamped at -70 mV to determine passive membrane properties and to examine spontaneous synaptic activity. Passive membrane properties were measured in voltage clamp mode. After breaking the seal, cell membrane properties (capacitance, input resistance, decay time constant) were determined by applying a 10 mV depolarizing step voltage command while holding the membrane potential at −70 mV and using the membrane test function integrated in pClamp 10.2 (Molecular Devices, Sunnyvale, CA). Spontaneous synaptic currents were recorded first in standard ACSF for 3-5 min at room temperature. At V_h_=-70 mV, in the absence of pharmacological blockers, the vast majority of synaptic currents are glutamatergic, with a minor contribution (about 15-25%) of GABAergic synaptic events. To examine glutamatergic currents in isolation (spontaneous excitatory postsynaptic currents, sEPSCs) BIC (10 µM), a GABA_A_ receptor antagonist, was added to the bath solution. GABA synaptic currents (spontaneous inhibitory postsynaptic currents, sIPSCs) were isolated by holding the membrane potential at +10 mV in ACSF, near the reversal potential for glutamate receptor-mediated synaptic currents. At this holding potential, glutamatergic synaptic currents are not present. Membrane currents were filtered at 1 kHz and digitized at 100-200 μs using Clampex 10.7 (gap-free mode). sPSCs were analyzed off-line (blinded to genotype) using the automatic detection protocol within the Mini Analysis Program (Synaptosoft) and subsequently checked manually for accuracy. The same program was used to calculate the average frequency, amplitude, area, rise time, and half-amplitude duration of all synaptic events recorded in a 90 sec period. Biocytin (0.2%) was added to the internal solution to obtain more detailed morphological data of recorded cells. After recordings, slices were fixed in 4% PFA and processed following standard procedures (Horikawa and Armstrong, 1988). DAB was used as the chromogen.

Additional recordings of CPNs were obtained in current clamp mode to complement observations made in voltage clamp mode and to examine action potential (AP) properties and intrinsic excitability. For current clamp recordings, the patch pipette contained (in mM): 112.5 potassium gluconate, 4 NaCl, 17.5 KCl, 0.5 CaCl_2_, 1 MgCl_2_, 5 ATP (dipotassium salt), 1 NaGTP, 5 EGTA, 10 HEPES. Cells were initially voltage clamped at -70 mV to determine passive membrane properties, then switched to current clamp mode to measure their resting membrane potential (RMP), AP amplitude, AP half-amplitude duration, rheobase, and intrinsic excitability. To measure rheobase, depolarizing current pulses (5 ms duration) of increasing intensity were injected until firing was achieved. Input-Output (I-O) functions were determined by counting the number of APs in response to 1 sec duration depolarizing current pulses of increasing intensity.

### Morphological measurements

Biocytin-labeled neurons were examined using a Zeiss confocal ApoTome microscope and images were acquired with the Zeiss Zen Blue software. Individual neurons were imaged at 10x, 40x and 100x (oil immersion) magnification, both as individual images, or as z-stack to acquire the full morphology of neuronal processes. Measurements of cell diameter, somatic area, and number of primary dendrites were obtained offline using the Zeiss Zen Blue software. Slices of the z-stack file were exported as individual TIFF files and then compiled in ImageJ, using the Z Project average intensity settings, into a single merged TIFF file. The biocytin illustrations in the figures were modified from the original black and white images acquired with Zen for pseudo-color and brightness/contrast enhancement using Adobe Photoshop. Further analysis (e.g., Sholl or spine density) was not attempted due to the variability of biocytin staining, the scarcity of spines in early development, and the slicing procedure itself, which generally makes the extent of dendritic fields difficult to determine.

### Statistical analyses

These experiments were done blindly as mouse genotypes were not identified until after weaning (at P21), ensuring scientific rigor and unbiased data analyses. Results are expressed as violin plots showing the median, lower and upper quartiles, as well as the distributions of numerical data points using density curves. Statistical analyses were performed using SigmaPlot (v.14) and data were analyzed using One-way ANOVAs (with Bonferroni correction) when 3 groups were compared. For comparisons between 2 groups we used the Student’s *t*-test or the Mann-Whitney U test. For differences in CPN excitability, Two-way RM ANOVA (with Bonferroni correction) was used. For comparisons between proportions, the Chi-square test was used. Differences between or among groups were deemed statistically significant if p < 0.05.

## Results

### Cerebral cortical development in WT and R6/2 mice

*Structural cortical abnormalities emerge during brain development in R6/2 transgenic mice:* As a preamble to the present study in the R6/2 mouse model of JHD, we first wanted to confirm our preliminary observations suggesting cytoarchitectural abnormalities in the cerebral cortex of symptomatic mice (Cepeda et al., 2019). In the previous study, about half of the R6/2 brains examined at >P60 days of age displayed abnormalities reminiscent of those observed in human FCD type 1, but not FCD type 2 or 3 according to the International League Against Epilepsy (ILAE) classification of FCDs (Abdijadid et al., 2015; Blumcke et al., 2011). These abnormalities included the blurring of cortical lamination, apical dendrite misorientation, neuronal disarray, and the occurrence of cortical zones devoid of the neuronal marker or, conversely, the agglutination of neurons in adjacent zones (Cepeda et al., 2019). In the present study, we extended our observations to both symptomatic (>P60) and pre-symptomatic R6/2 (<P35) mice, in an attempt to differentiate neurodegeneration from cortical maldevelopment. The brains of WT and R6/2 mice at P14 (n=5, WT; n=5, R6/2), P21 (n=2, WT; n=3, R6/2), P30 (n=4, WT; n=5, R6/2) and P60 (n=3 WT; n=4, R6/2) were perfused, fixed, and immunostained for the specific neuronal marker NeuN. Confirming and extending previous observations, we found signs of FCD not only in symptomatic but also in pre-symptomatic HD brains **(Fig. 1)**. At P21, 3/6 (50%) samples were positive for FCD-like areas (2 were in motor and 1 in somatosensory cortex); at P30, 5/7 (71%) were positive (3 in motor and 2 in somatosensory cortex), and at P60, 5/6 (83%) samples showed signs of dysplasia (3 in motor and 2 in somatosensory cortex). Thus, about 68% of cortical samples within the age ranges examined displayed architectural abnormalities. At P14, dysplastic cortical lesions were more subtle or non-existent (only 2 out of 10 samples showed very mild FCD-like features). None of the WT samples showed signs of FCD.

**Fig. 1:**
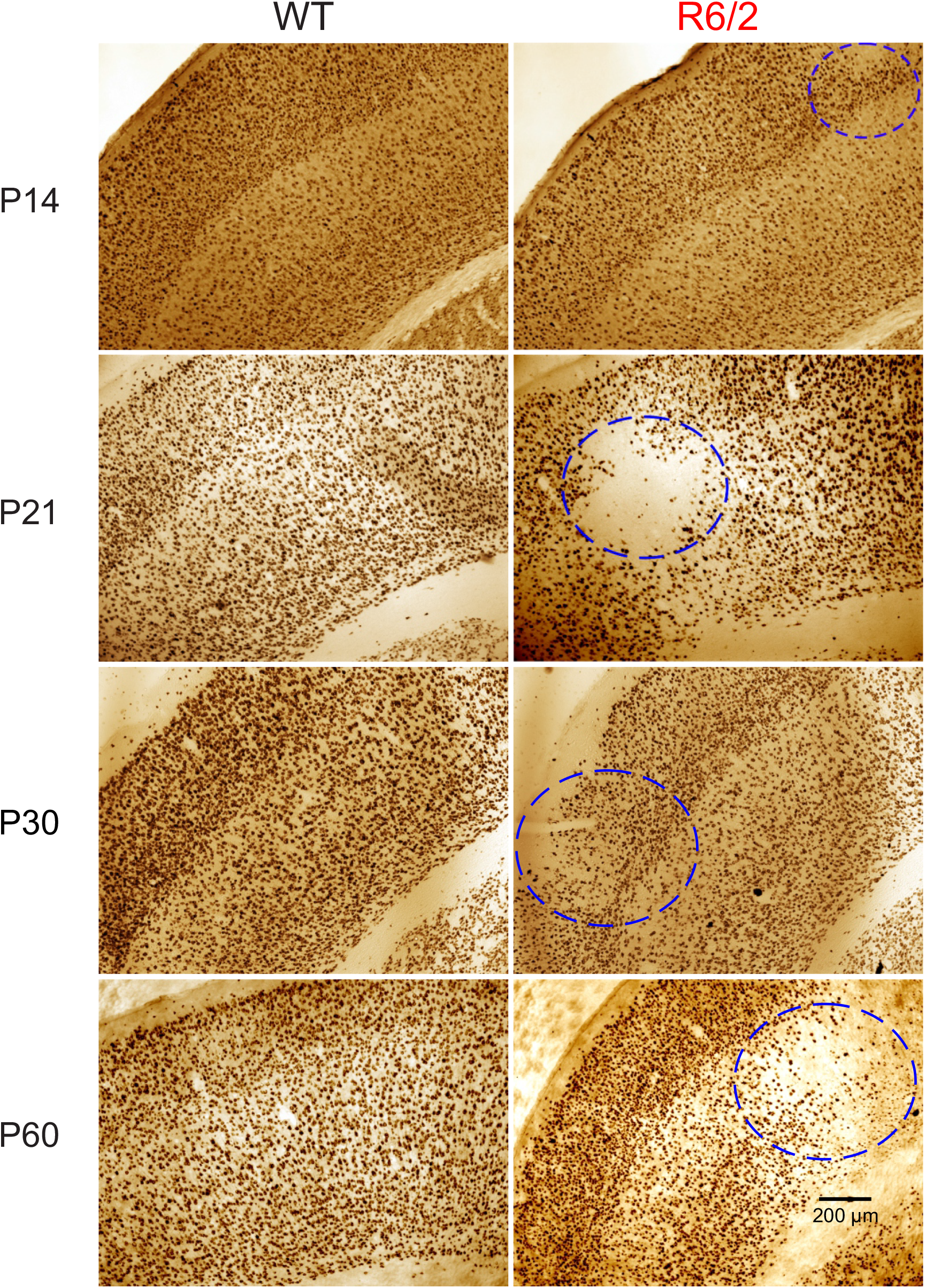
NeuN immunohistochemistry in R6/2 sensorimotor cortex. NeuN, a specific neuronal marker was used to examine the cytoarchitecture of the sensory and motor cortices of WT and R6/2 mice. NeuN staining demonstrated architectural abnormalities in developing R6/2 mice. Very subtle at P14, they became much more pronounced at P21, P30, and P60. The panels show hemi- slices (30 µm thick) of sensorimotor cortex from P14-P60 animals immunostained with NeuN. Circles on the right panels indicate the areas with highest abnormality. Calibration applies to all panels.

Overall, the findings of cytoarchitectural alterations in the cortex of developing HD mice confirmed previous findings and prompted us to undertake a detailed examination and comparison of cortical and striatal neuron development in R6/2 mice compared to age-matched WT controls.

Our main focus was on the developmental profile of intrinsic passive and active cell membrane properties and spontaneous synaptic currents of CPNs and MSNs.

*The intrinsic membrane properties of CPNs are altered in developing R6/2 mice:* Biophysical membrane properties [cell capacitance (Cm), input resistance (Rm), and decay time constant (tau)], provide relevant information and can be used as proxies for the extent of cell surface membrane area (Cm) and density of functional ion channels (Rm). **Fig. 2A** shows membrane properties of WT and R6/2 CPNs at P7, P14 and P21 recorded in voltage clamp mode (V_h_=-70 mV) using Cs-methanesulfonate as the patch pipette internal solution. We first compared genotype differences in these measures within each age group using the non-parametric Mann-Whitney test. These comparisons demonstrated that cell membrane capacitance was significantly reduced in R6/2 CPNs at P14 and the input resistance was significantly increased both at P7 and P14 but not at P21. As expected, decreased cell capacitance at P14 was accompanied by a significant reduction in decay time constant. Overall, these findings underscore that very early in development (P7 and P14), CPNs from R6/2 mice display reduced membrane area, as well as reduced number of functional channels manifested by significant increases in input resistance. This suggests that maturation of CPNs in R6/2 mice is delayed compared to that of WT neurons.

**Fig. 2:**
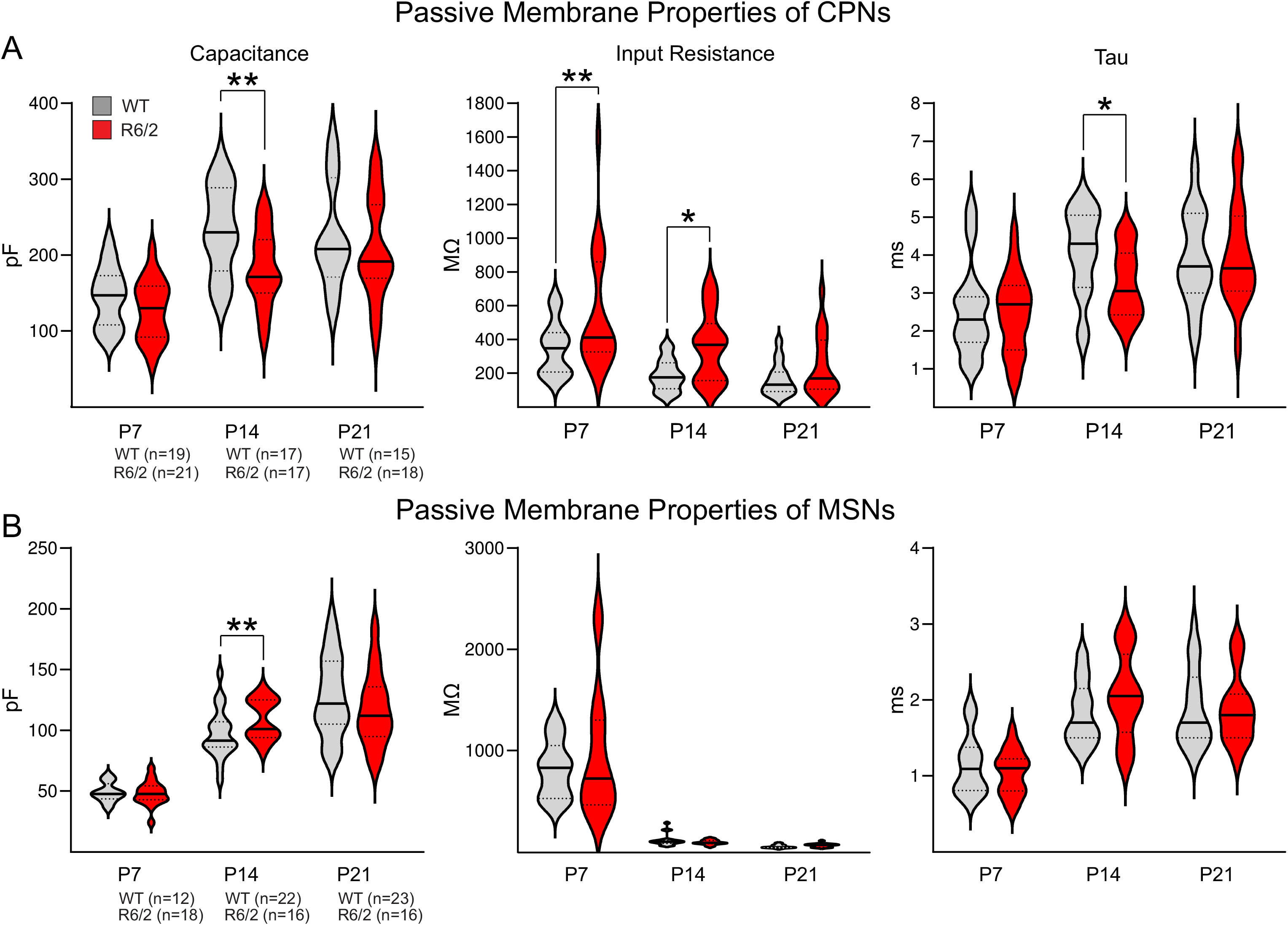
Passive membrane properties of CPNs and MSNs. Passive membrane properties (capacitance, input resistance, and decay time constant of CPNs **(A)** and MSNs **(B)** recorded in voltage clamp mode and with Cs-methanesulfonate in the internal pipette solution. Violin plots show the median (solid line), lower and upper quartiles (dotted lines), as well as the distributions of numerical data points using density curves. **(A)** Cell membrane capacitance (in pF), input resistance (in MΩ), and decay time constant (Tau, in ms) in WT and R6/2 CPNs at P7, P14, and P21. Numbers in parentheses indicate the number of cells recorded in WT and R6/2 mice for each age. **(B)** Cell membrane properties of MSNs. In this and subsequent figures, asterisks indicate statistically significant differences; * p<0.05, ** p<0.01, and *** p<0.001, as determined by Mann-Whitney U test or unpaired Student’s t-test.

To examine the developmental trajectories of passive membrane properties of WT and R6/2 CPNs we used One-way ANOVAs followed by pairwise multiple comparisons (Bonferroni *post hoc* t-tests). For cell membrane capacitance, we found a statistically significant age-dependent effect (p<0.001, One-way ANOVA) for WT and R6/2 CPNs. Bonferroni t-tests demonstrated that CPNs membrane capacitance was significantly smaller at P7 compared to both P14 (WT, p<0.001; R6/2, p=0.007) and P21 (WT, p<0.001; R6/2, p<0.001), but not between P14 and P21 **(Suppl. Fig. 1)**. Reduced membrane area is generally associated with increased input resistance. In agreement, there was a statistically significant age effect (p<0.001) in both WT and R6/2 CPNs. Pairwise multiple comparisons demonstrated significant differences in cell membrane input resistance between P7 and both P14 (p<0.001) and P21 (p<0.001) in WT CPNs, but not between P14 and P21. In R6/2 CPNs, the difference between P7 and P21 was statistically significant (p=0.002), but between P7 and P14 there was just a non-significant trend (p=0.076). With regard to the decay time constant, there also was a significant effect among the different ages (p<0.001). The time constant was significantly faster at P7 compared to both P14 (WT, p=0.001; R6/2, p=0.038) and P21 (WT, p=0.006; R6/2, p<0.001) but not between P14 and P21. These observations indicate that the main increase in cell membrane area and density of functional channels in both WT and R6/2 mice occurs between P7 and P14, albeit the decrease in input resistance of R6/2 CPNs was not as dramatic as in WTs.

For the most part, observations obtained with Cs-methanesulfonate were confirmed when recordings were obtained in voltage clamp mode using K-gluconate, i.e., in general the cell membrane capacitance was reduced while the input resistance was increased. In particular, at P14 the reduction in R6/2 CPN capacitance compared to WT was statistically significant (p=0.011) and the difference persisted at P21 (p<0.001). Changes in the RMP were examined in current clamp mode. There were no significant differences between genotypes at any age (Table I). At P14, R6/2 CPNs were more depolarized than WT CPNs, but the difference did not reach statistical significance (p=0.086). As for age-dependent effects, the main change in RMP was from P7 to P14, as CPNs became more hyperpolarized. The age effect was highly significant (p<0.001). No further changes in RMP occurred between P14 and P21 and this was true for both WT and R6/2 CPNs. AP properties and CPN excitability also were examined in current clamp mode. There were no differences in AP amplitude, threshold for firing, and AP half-amplitude duration between genotypes in any age group. Interestingly, the rheobase (i.e., the minimum current intensity needed to trigger an AP) was significantly reduced in R6/2 CPNs compared to WTs at P14 (Table I **and** **Fig. 3B)**. We thus examined CPN excitability by constructing I-O functions. We found that I-O functions were similar between WT and R6/2 CPNs at P7 and P21. In contrast, at P14 R6/2 CPNs were significantly more excitable than WT CPNs **(Fig. 3C)**. This may be explained by the increased input resistance of R6/2 CPNs at this age, reduced rheobase, and a slightly more depolarized RMP of CPN from R6/2 mice compared to WTs.

**Fig. 3:**
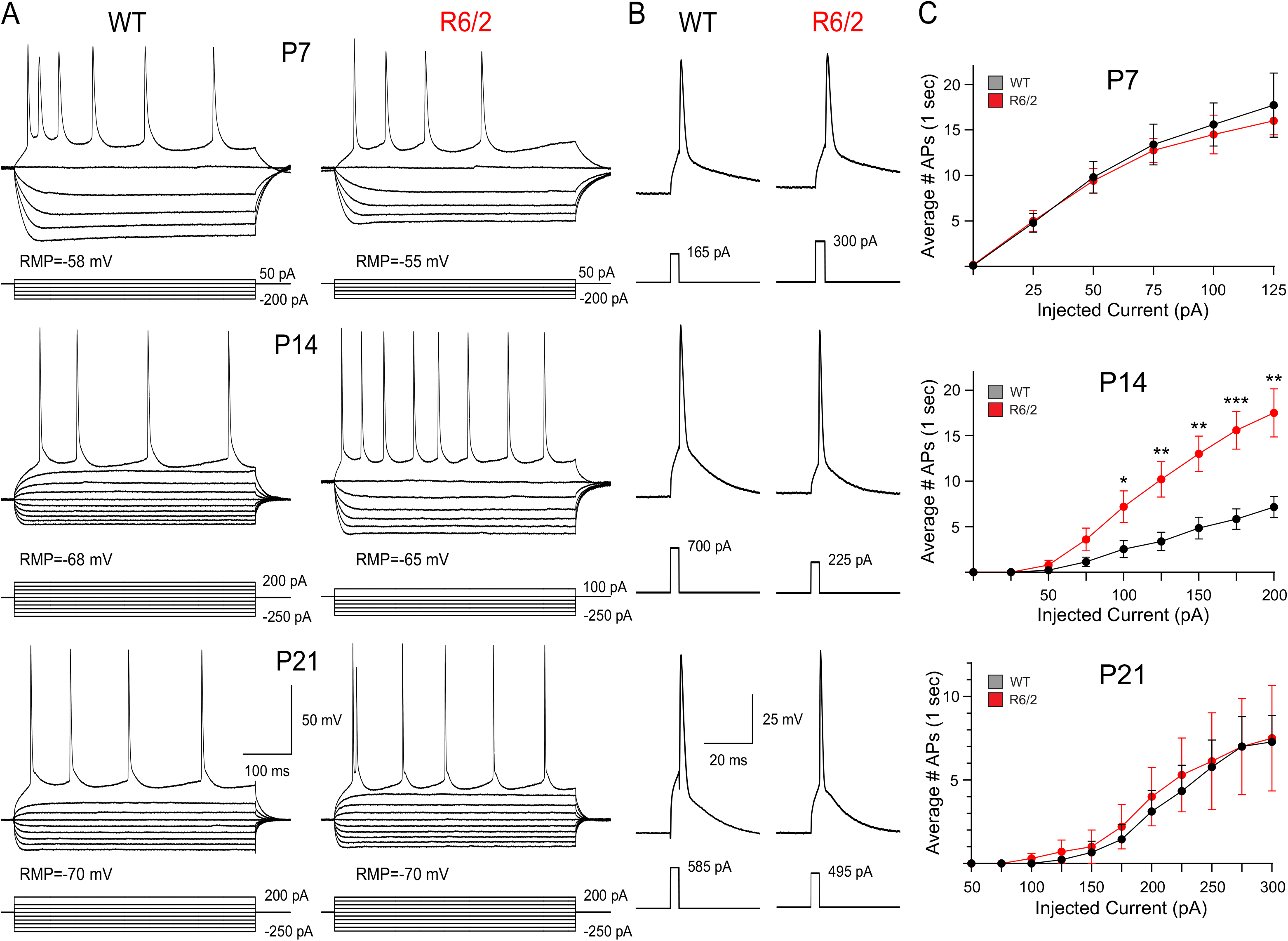
Cortical excitability during development. **(A)** Voltage changes induced by current injection of hyperpolarizing and depolarizing current pulses (500 ms duration) in WT and R6/2 CPNs at P7, P14, and P21. Notice the higher number of APs in R6/2 CPNs at P14, even with a lower intensity of the positive current. **(B)** Minimum current required to induce an AP at WT and R6/2 CPNs. Rheobase was consistently lower in R6/2 CPNs at P14. **(C)** Input-output functions demonstrated statistically significant increases in R6/2 CPN excitability compared to WTs at P14 (Two-way RM ANOVA).

*Spontaneous synaptic activity is reduced early in R6/2 CPNs but is similar to WT values at P21:* The primary source of glutamatergic inputs to CPNs is from thalamo-cortical and intra- telencephalic CPN terminals (Thomson and Lamy, 2007). GABAergic inputs are provided by local circuit interneurons, in particular from parvalbumin (PV)- and somatostatin (SOM)-expressing interneurons. We examined the development of spontaneous glutamatergic and GABAergic synaptic activity of CPNs in voltage clamp mode with Cs-methanesulfonate as the internal patch pipette solution.

Differences between WT and R6/2 CPNs were compared using the Mann-Whitney test **(Fig. 4)**. At V_h_=-70 mV in ACSF, a statistically significant deficit of synaptic inputs occurred in R6/2 compared to WT CPNs at P7 and P14. At P21 there was no statistically significant difference between genotypes. After isolation of glutamatergic synaptic currents with BIC, the reduction in sEPSC frequency at P7 persisted. In contrast, at P14 and P21, the frequency was similar to WT levels. GABAergic synaptic activity also was significantly reduced in CPNs from R6/2 mice at P7. At P14, sIPSC frequency remained reduced but the difference was not statistically significant and at P21 no significant differences occurred. These findings indicate that, during early development, synaptic inputs in R6/2 CPNs lag behind those in WTs. Later in development, inputs to CPNs in R6/2 mice compensate and are similar in frequency to WTs.

**Fig. 4:**
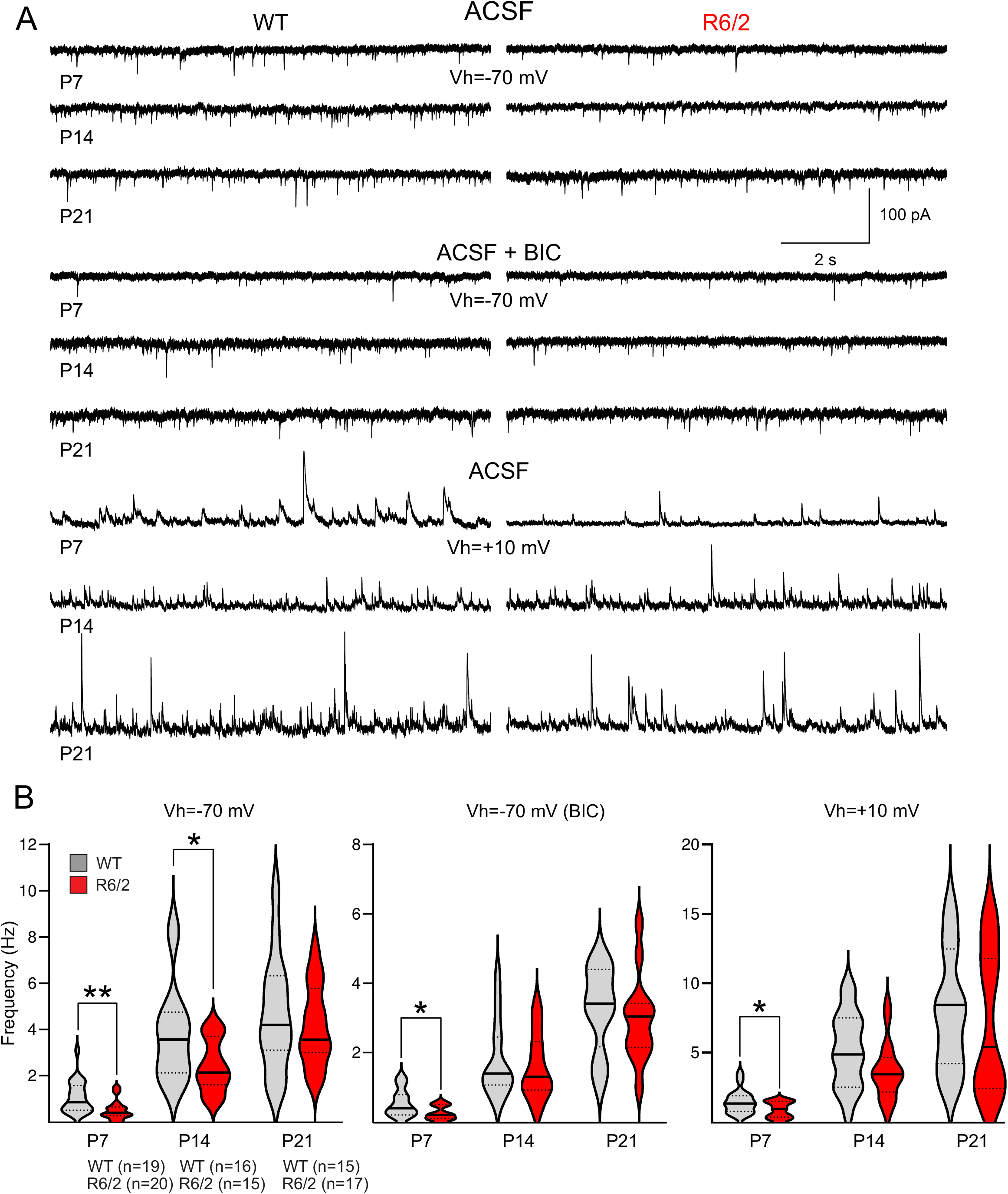
**Development of spontaneous synaptic activity of CPNs in WT and R6/2 mice**. **(A)** Traces are representative recordings (10 sec duration) of spontaneous synaptic events in CPNs from WT and R6/2 mice a P7, P14, and P21. Top traces show synaptic activity in ACSF, middle traces are isolated glutamate events after addition of BIC, and bottom traces are GABAergic events in ACSF after stepping the membrane to +10 mV, close to the glutamate reversal potential. **(B)** Violin plots show the median (solid line), lower and upper quartiles (dotted lines), as well as the distributions of numerical data points using density curves for WT an R6/2 CPNs at P7, P14, and P21. In all cases, the medians in R6/2 CPNs were lower than those in WTs, and at P7 the reduced inputs were statistically significant. Numbers in parentheses indicate the number of cells recorded in WT and R6/2 mice at each age.

The developmental trajectories of synaptic activity in WT and R6/2 CPNs were analyzed by One-way ANOVA followed by Bonferroni t-tests. At V_h_=-70 mV in ACSF with no blockers added, we found a statistically significant age effect (p<0.001) for both WT and R6/2 CPNs **(Suppl. Fig. 2)**. With development, spontaneous synaptic currents increased in frequency. In WT mice, significant differences were observed between P7 and both P14 (p<0.001) and P21 (p<0.001), but not between P14 and P21 (p=0.491), suggesting that by P14 CPNs from WT mice already receive a nearly full complement of synaptic inputs. For R6/2 mice, significant differences were seen not only between P7 and both P14 (p<0.001) and P21 (p<0.001), but also between P14 and P21 (p<0.001). One possible interpretation is that the development of synaptic inputs onto CPNs of WT and R6/2 mice follow different trajectories and confirm the above assumption that the full complement of synaptic inputs reaching R6/2 CPNs is delayed. At V_h_=-70 mV in the presence of BIC (i.e., isolated glutamatergic inputs), a significant age effect (p<0.001) also occurred in both WT and R6/2 mice. Bonferroni t-tests demonstrated significant differences between P7 and both P14 (WT, p=0.032; R6/2, p<0.001) and P21 (WT, p<0.001; R6/2, p<0.001), and also between P14 and P21 (WT, p=0.024; R6/2, p<0.001). At V_h_=+10 mV (i.e., GABAergic inputs), a significant age effect occurred (p<0.001) in both WT and R6/2 mice. Pairwise multiple comparisons demonstrated significant differences between P7 and both P14 (WT, p=0.001; R6/2, p=0.027) and P21 (WT, p<0.001; R6/2, p<0.001), and also between P14 and P21 (WT, p=0.023; R6/2, p=0.004).

*Changes in spontaneous synaptic currents amplitude, area, and kinetics during development:* We measured potential differences in amplitude, area, and some kinetic properties (rise time and half-amplitude duration) of glutamatergic and GABAergic synaptic events between genotypes and throughout cortical development **(Suppl. Table II)**. In general, no significant genotype effects were found for any of these properties. The only significant difference was an increase in the area (p=0.024, Mann-Whitney test) and rise time (p=0.021) of glutamatergic synaptic events (ACSF + BIC) in R6/2 compared to WT CPNs at P14. However, statistically significant age-dependent changes were observed regardless of genotype. The amplitude and area of synaptic events recorded at V_h_=-70 mV in ACSF (with or without BIC) and at V_h_=+10 mV increased progressively from P7 through P21. Changes in rise time were less evident and more inconsistent. The most conspicuous change in kinetics was a statistically significant reduction in half-amplitude duration of GABAergic events from P7 to P14 in both WT and R6/2 CPNs, probably due to changes in subunit composition of GABA_A_ receptors.

*The presence of Giant Depolarizing Potentials in R6/2 CPNs is increased compared to that in WTs:* A characteristic signature of immature cortical networks is the occurrence of Giant Depolarizing Potentials (GDPs) in projection neurons (Ben-Ari et al., 1989). In rodents, this activity progressively disappears around P7 and it is not observed at P14 or later. GDPs are generated by the interplay among GABA_A_ receptors, glutamate receptors, and voltage-gated Ca^2+^ channels (Ben-Ari et al., 1997; Leinekugel et al., 1997). The principal generator of GDPs is the synchronous release of GABA from PV- and SOM-expressing interneurons (Wester and McBain, 2016), as blockers of GABA_A_ receptors completely abolish GDPs (Ben-Ari et al., 1989). Thus far, there are no studies examining the development of GDPs in CPNs of HD model mice. We examined the occurrence of GDPs in WT and R6/2 CPNs at P7 using Cs-methanesulfonate internal solution at V_h_=-70 mV and at V_h_=+10 mV. In voltage clamp mode, GDPs are manifested by very large inward (at -70 mV) or outward (at +10 mV) currents with a typical V-shape. They are easily identified by their rhythmicity, significantly larger amplitude (>100 pA at V_h_=-70 mV and >400 pA at V_h_=+10 mV) and longer duration (half-amplitude duration >100 ms) compared to sEPSCs and sIPSCs. To determine the reversal potential for GABA currents we stepped the voltage command (from V_h_=-70 mV) by increments of 20 mV in the depolarizing direction. We found the giant GABA current reversed at ∼-57 mV **(Fig. 5)**. Bath application of BIC abolished GDPs completely (**Suppl. Fig. 3F).** Interestingly, in current clamp mode, some GDPs evoked action potentials as a result of the membrane depolarization reaching the threshold for firing **(Suppl. Fig. 3**E-F).

**Fig. 5:**
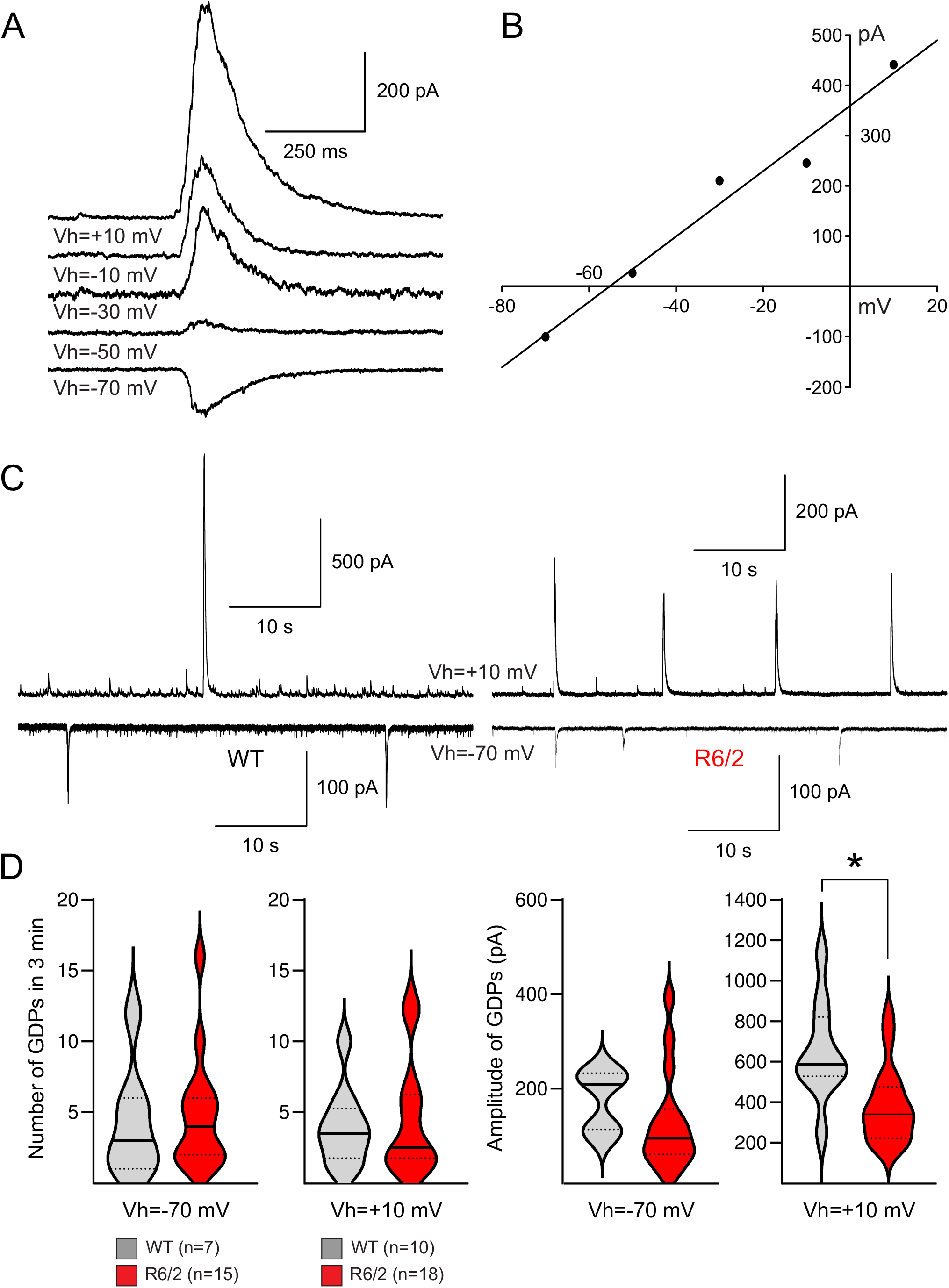
**GDPs recorded in CPNs of WT and R6/2 mice at P7**. **(A)** To determine the GABA current reversal potential, a CPN displaying spontaneous GDPs was stepped to more depolarized membrane potentials from a holding voltage of -70 mV in 20 mV increments. At -50 mV, the outward current was very small, but as the membrane was depolarized the amplitude of the current increased. **(B)** The graph shows the IV (current-voltage) relationship of the GABA current. The equilibrium potential was ∼-57 mV. **(C)** Spontaneous GDPs recorded at -70 and +10 mV holding potentials in WT and R6/2 CPNs. **(D)** Violin plots showing the median (solid line), lower and upper quartiles (dotted lines), of the number of GDPs in 3 min recordings, as well as the amplitude of GDPs. No significant differences were observed in the number of GDPs at either -70 mV or +10 mV holding potentials. However, the amplitude of GDPs was reduced at both holding potentials and the difference was statistically significant at +10 mV.

At -70 mV holding potential, more R6/2 CPNs displayed GDPs compared to WTs (71.4% R6/2 vs. 36.8% WT, n=19 and 21 CPNs respectively, p=0.028, Chi-square). Similarly, at V_h_=+10 mV, the percent of CPNs displaying GDPs (85.7% R6/2 vs. 52.6% WT) was significantly higher (p=0.023, Chi-square). The fact that more cells from R6/2 mice displayed GDPs further supports the idea that CPNs at P7 are less mature in R6/2 compared to WT mice. We also estimated the relative frequency of GDPs during the 3 min recording period. In the subset of CPNs displaying GDPs, the frequency was similar in WT and R6/2 CPNs (at V_h_=-70 mV, WT, 4.3±1.5, and R6/2, 4.8±1; at V_h_=+10 mV, WT, 3.9±0.85 and R6/2, 4.6±0.95). Interestingly, the amplitude of GDPs was reduced in R6/2 compared to WT CPNs. While at V_h_=-70 mV, the amplitude of GDPs in WT *vs.* R6/2 CPNs was not statistically significant (p=0.162, Mann-Whitney test), the difference between WT and R6/2 CPNs recorded at V_h_=+10 mV reached statistical significance (p=0.004). *Paroxysmal Discharges in CPNs from WT and R6/2 mice:* A typical feature of JHD is the occurrence of epileptic seizures. They also are common in the R6/2 model of HD after the onset of motor symptoms (between 5-7 weeks of age). The cause of seizures is likely an imbalance between excitation and inhibition in the cerebral cortex (Cummings et al., 2009; Spampanato et al., 2008). *In vitro* bath application of BIC induces epileptiform paroxysmal discharges (PDs) due to generalized cortical disinhibition (Cummings et al., 2009). We examined the effects of BIC application (10 µM) in WT and R6/2 CPNs during brain development. The presence of PDs after BIC was determined by three main parameters; duration of the PD (>400 ms), amplitude of the PD (generally >1500 pA), and classification of PDs as simple or complex (one vs. multiple components). At P7, 69% of WT CPNs displayed PDs and in R6/2 CPNs, 67% of neurons displayed PDs. The difference in distribution was not statistically significant (p=0.89, Chi-square test). At P14, the ability to generate epileptiform activity increased considerably, likely due to higher abundance of excitatory inputs. All CPNs, regardless of genotype, presented with PDs after BIC blockade of GABA_A_ receptors. At P21, almost all CPNs from R6/2 mice presented with PDs (11/12, 91.7%), whereas in WTs slightly fewer cells displayed PDs (8/11, 72.7%, Chi square, p=0.23). We further compared the frequency and duration of PDs in CPNs recorded from R6/2 and WT littermates on the same recording day and regardless of age (i.e., data from P7, P14, and P21 pooled). These side-by-side comparisons revealed some differences in PD duration and complexity **(Suppl. Fig. 4)**. While the average frequency of PDs in 8 min BIC application was not different, the duration of PDs was significantly increased in R6/2 CPNs (p=0.049, Student’s t-test). *NMDA spikes:* NMDA spikes are regenerative dendritic depolarizations produced by local activation of NMDA receptors and were first recorded in the basal dendrites of CPNs (Schiller et al., 2000). In general, they are evoked by focal glutamate uncaging or stimulation of glutamatergic afferents, although very rarely they may occur spontaneously (Antic et al., 2010; Kumar et al., 2018; Oikonomou et al., 2015). They can be easily recognized by their peculiar morphology, relatively long duration, and higher amplitude compared to regular spontaneous synaptic events. Occasionally, they also are associated with Na^+^ spikelets. We recorded a number of these dendritic NMDA spikes in CPNs from both WT and R6/2 mice at P14 and P21, but not at P7 **(Fig. 6A)**. We could measure 4 NMDA spikes in WT and 8 in R6/2 CPNs (over both ages). The mean amplitude was 123.1±33.9 pA in WT and 152.4±44.2 pA in R6/2 (p=0.676, Student’s *t*-test). The duration was 560.7±129.9 ms in WT and 781±111.1 ms in R6/2 (p=0.255) and the half-amplitude duration was 232.6±42.2 ms in WT and 408.8±33.984.9 ms in R6/2 (p=0.193).

**Fig. 6:**
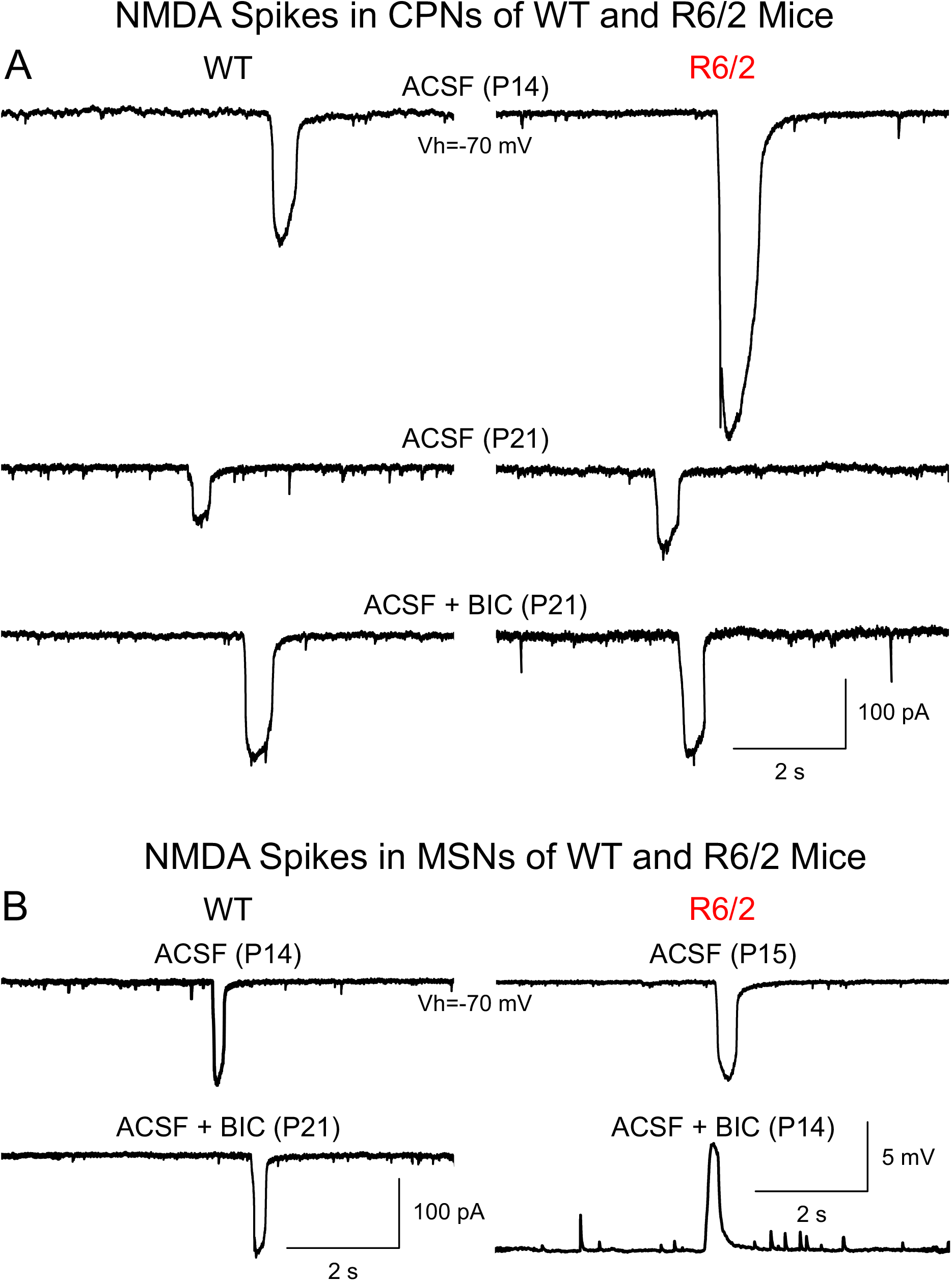
NMDA spikes. Spontaneous NMDA spikes recorded in CPNs **(A)** and MSNs **(B)** of WT and R6/2 mice aged P14 and P21. Although NMDA spikes occur very rarely, a few NMDA spikes were observed under both ACSF and ACSF + BIC. Their morphology is very typical and can be differentiated by their large amplitude and duration compared with other spontaneous synaptic currents. All traces were recorded in voltage clamp mode (V_h_=-70 mV), except for the last one (R6/2 MSN), which was recorded in current clamp mode.

*Morphological abnormalities of CPNs revealed by biocytin labeling:* As mentioned in a preliminary communication (Cepeda et al., 2019), IR-DIC microscopy and biocytin staining provided evidence of CPN misorientation and the presence of dysmorphic processes typically seen in pediatric FCD (Cepeda et al., 2003b). This was confirmed in the present study. First we selected the biocytin-stained CPNs with the most intact morphology from P7-P21 (P7, WT, n=11, R6/2, n=9; P14, WT, n=5, R6/2, n=11; P21, WT, n=5, R6/2, n=8). We found a number of R6/2 CPNs displaying dysmorphic processes including tortuous dendrites and/or axons, sharp bends (about 90°), and misoriented apical dendrites. Only 1 WT CPN (5%) at P7 showed a meandering apical dendrite, whereas 10 R6/2 CPNs (38.5%) displayed such abnormality (3/8 at P7, 4/10 at P14, and 3/8 at P21) **(Fig. 7)**. The difference between genotypes was statistically significant (p=0.0105, Chi-square). In addition, one of the CPNs also showed a tortuous axon **(Suppl. Fig. 5B1)**, while another had double apical dendrites **(Suppl. Fig. 5C**), and yet another CPN was misoriented **(Suppl. Fig. 5E-E1)**. One cell with tortuous apical dendrite at P7 also showed sinuosity of the terminal tuft in layer I **(Suppl. Fig. 5A-A2)**.

**Fig. 7:**
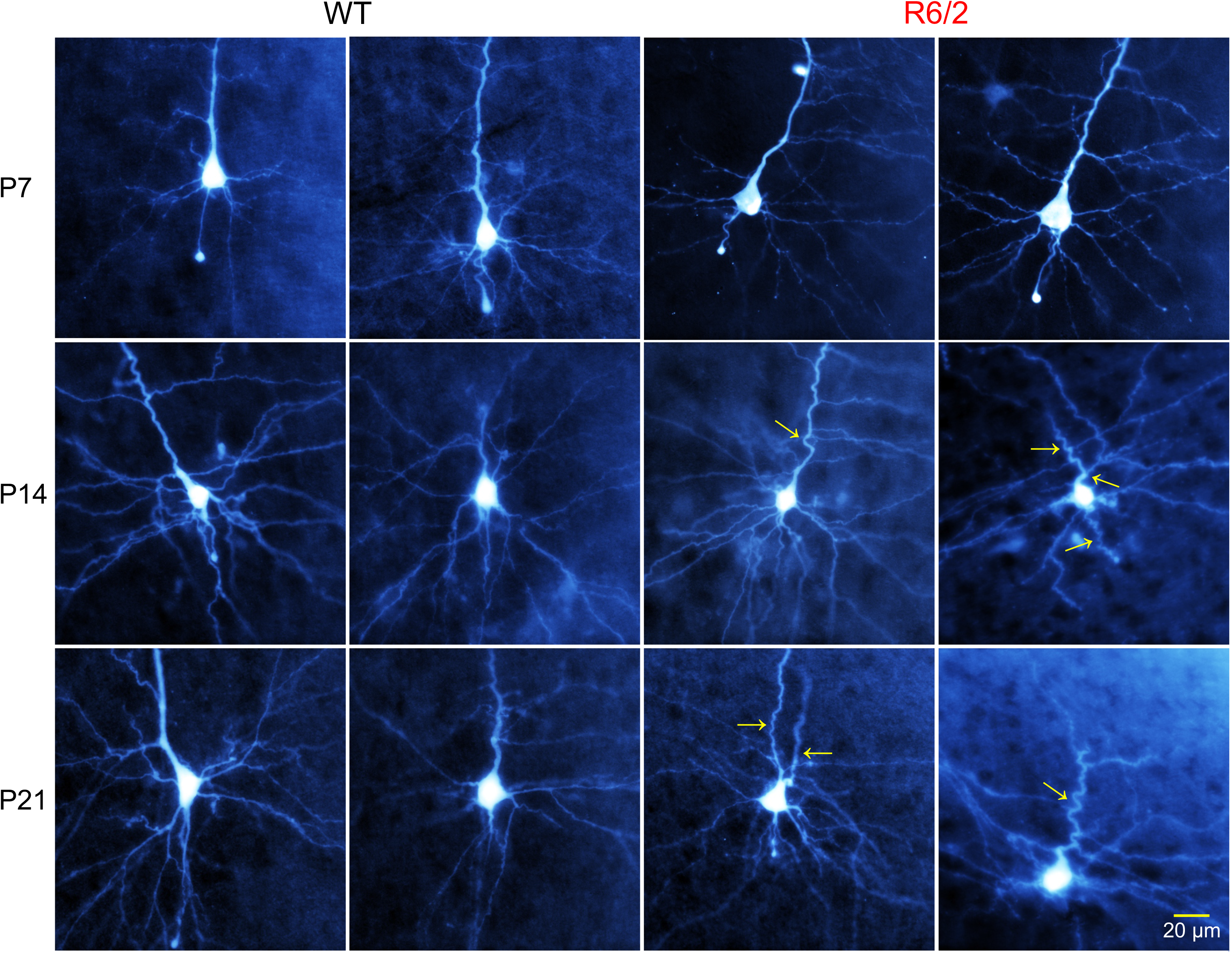
Morphology of CPNs from WT and R6/2 mice at P7, P14, and P21. CPNs were recorded and filled with biocytin. Sharp bends of the apical dendrites and tortuous processes were frequently observed in R6/2 CPNs (yellow arrows), whereas in WT neurons this occurred only rarely. One R6/2 CPN at P21 had two apical dendrites. The original black and white images acquired with Zen were modified for pseudo-color and brightness/contrast enhancement using Adobe Photoshop. Calibration in the bottom right applies to all panels.

We also measured somatic area, diameter, and number of primary dendrites of WT and R6/2 CPNs. For somatic area, no differences were found at P7 (WT, 146.1±6.95 µm^2^; R6/2, 153.4±18.8 µm^2^, p=0.691, n=11 and 9 respectively) and P14 (WT, 192.9±33.5 µm^2^; R6/2, 171.3±15.3 µm^2^, p=0.526, n=5 and 11 respectively). However, at P21 a trend for reduced somatic area was found (WT, 198.3±21.9 µm^2^; R6/2, 152.1±10.6 µm^2^, p=0.056, n=4 and 8 respectively).

For cell diameter, no statistically significant differences were found (P7 WT, 14.4±0.46 µm; R6/2, 14.9±1.1 µm, p=0.656; P14 WT, 17.04±1.6 µm; R6/2, 15.5±0.7 µm, p=0.322; P21 WT, 17.0±1.1 µm; R6/2, 15.3±0.6 µm, p=0.0631). Similarly, no difference in the number of primary dendrites (in general 4-5) between genotypes was observed.

## Striatal development in WT and R6/2 mice

*Membrane properties of striatal MSNs:* In the same mice used to examine CPNs we measured membrane properties of MSNs in WT and R6/2 mice at P7, P14 and P21 **(Fig. 2B)**. Overall, no major differences were observed in the membrane properties of WT and R6/2 MSNs, except for an unexpected significant increase in cell capacitance of R6/2 MSNs compared to WT MSNs at P14 (p=0.037, Mann-Whitney test). This is in contrast to CPNs, which in general have reduced membrane capacitance throughout development, and suggest early but transient compensatory mechanisms in striatal MSNs. There were significant age-dependent effects in WT and R6/2 MSNs **(Suppl. Fig. 1)**. For cell membrane capacitance, we found a statistically significant effect among different ages (p<0.001). Bonferroni t-test demonstrated significant differences in WT MSNs capacitance between P7 and both P14 (p<0.001) and P21 (p<0.001). There also was a significant difference between P14 and P21 (P<0.001). These observations indicate that, with development, there is a steady increase in cell capacitance of WT MSNs from P7 until P21. In contrast, in R6/2 MSNs, while there was a significant increase in cell capacitance between P7 and both P14 (p<0.001) and P21 (p<0.001), the difference between P14 and P21 was not statistically significant. This could indicate that in R6/2 MSNs membrane area stops increasing by P14. In terms of changes in cell membrane input resistance, there was a statistically significant effect among different ages (p<0.001) in both WT and R6/2 MSNs. Bonferroni t-test demonstrated significant differences in cell membrane input resistance between P7 and both P14 (WT, p<0.001; R6/2, p<0.001) and P21 (WT, p<0.001; R6/2, p<0.001) MSNs, but not between P14 and P21 (WT, p=0.298; R6/2, p=1.00).

With regard to the decay time constant, there also was a significant age effect (p<0.001) in both WT and R6/2 neurons. The time constant was significantly shorter at P7 compared to both P14 (WT, p<0.001; R6/2, p<0.001) and P21 (WT, p<0.001; R6/2, p<0.001), but not between P14 and P21 (WT, p=1.00; R6/2, p=0.876).

*Development of spontaneous synaptic activity in MSNs:* The main source of glutamatergic inputs onto MSNs is from cortical (Bolam et al., 2000) and thalamic (CM/Pf nuclear complex) terminals (Smith et al., 2014). GABAergic inputs are provided by local circuit interneurons, in particular from PV- and SOM-expressing interneurons (Tepper et al., 2018) and a small contribution from cortical SOM interneurons (Rock et al., 2016). In the same mice used for recordings of CPNs we also examined, in parallel, the development of spontaneous glutamatergic and GABAergic synaptic activity impinging on MSNs under similar recording conditions, In ACSF, at V_h_=-70 mV, the frequency of spontaneous synaptic currents in R6/2 MSNs, compared to WTs, decreased with age **(Fig. 8)**. Bonferroni t-test demonstrated significant genotype differences at P21 (p=0.008), but not at P7 (p=0.865) or P14 (p=0.712). Similar findings were observed for frequency of synaptic activity in the presence of BIC. At P7 or P14, no genotype differences occurred (p=0.888 and p=0.514 respectively), however, at P21 there was a significant reduction in glutamatergic inputs to R6/2 MSNs compared to WTs (p=0.004). For changes in frequency of GABAergic synaptic activity, we found no significant genotype differences at P7 (p=0.846) or P21 (p=0.10). Interestingly, there was a transient significant decrease of GABAergic inputs to R6/2 MSNs compared to WTs at P14 (p=0.009).

**Fig. 8:**
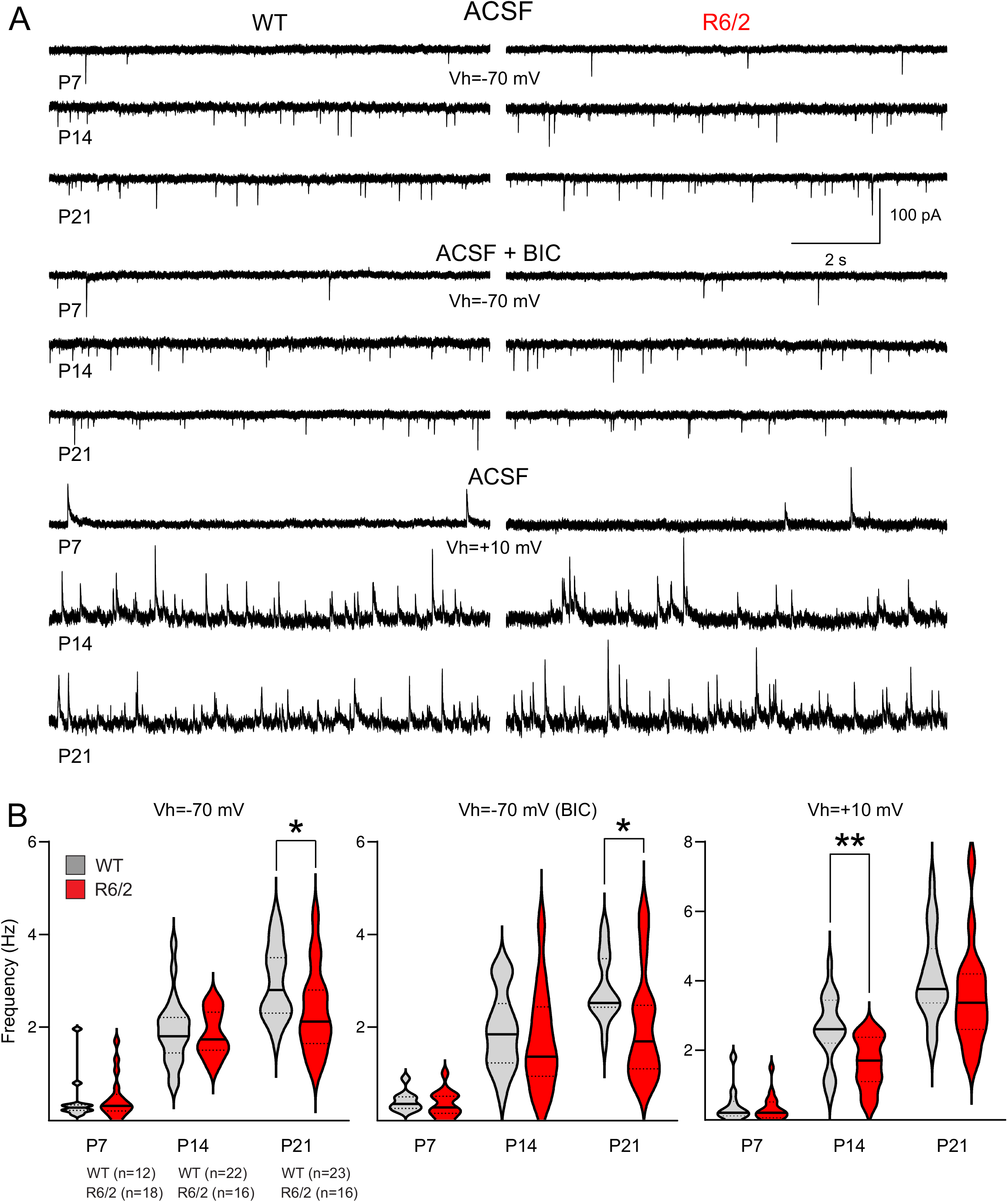
Development of spontaneous synaptic activity of MSNs in WT and R6/2 mice. **(A)** Traces are representative recordings (10 sec duration) of spontaneous synaptic events in MSNs from WT and R6/2 mice a P7, P14, and P21. Top traces show synaptic activity in ACSF, middle traces are isolated glutamate events after addition of BIC, and bottom traces are GABAergic events in ACSF after stepping the membrane to +10 mV. **(B)** Violin plots show the median (solid line), lower and upper quartiles (dotted lines), as well as the distributions of numerical data points using density curves for WT an R6/2 CPNs at P7, P14, and P21. At -70 mV, with or without BIC, the spontaneous synaptic activity was significantly reduced at P21. The frequency of GABAergic currents was significantly reduced at P14 but recovered at P21. Numbers in parentheses indicate the number of cells recorded in WT and R6/2 mice for each age.

At V_h_=-70 mV in ACSF with no blockers added, there was a statistically significant age-dependent effect (p<0.001) in both WT and R6/2 MSNs **(Suppl. Fig. 2)**. Spontaneous synaptic currents in MSNs from both WT and R6/2 mice increased in frequency with age. In WT mice, Bonferroni t-tests demonstrated significant differences between P7 and both P14 (p<0.001) and P21 (p<0.001), and also between P14 and P21 (p<0.001), suggesting a steady increase in the frequency of synaptic events during the phase of development. A slightly different picture emerged in R6/2 mice. While statistically significant increases in synaptic inputs occurred between P7 and both P14 (p<0.001) and P21 (p<0.001), the increase between P14 and P21 was not significant (p=0.087). At V_h_=-70 mV in the presence of BIC, a significant age effect also occurred (p<0.001). In WT mice, Bonferroni t-tests demonstrated significant differences between P7 and both P14 (p<0.001) and P21 (p<0.001), and also between P14 and P21 (p<0.001). In contrast, in R6/2 MSNs, while the differences between P7 and both P14 (p=0.002) and P21 (p<0.001) were highly significant, the difference between P14 and P21 was not statistically significant. This implies that WT MSNs display a gradual increment in glutamatergic input frequency during early brain development, whereas the inputs in R6/2 MSNs remain stationary from P14 to P21. At V_h_=+10 mV (i.e., GABAergic inputs), significant age effects were observed (p<0.001) in both WT and R6/2 mice. Bonferroni t-tests demonstrated significant differences between P7 and both P14 (WT, p<0.001; R6/2, p=0.002) and P21 (WT, p<0.001; R6/2, p<0.001), and also between P14 and P21 (WT, p<0.001; R6/2, p<0.001).

We also measured potential differences in amplitude, area, and some kinetic properties of glutamatergic and GABAergic synaptic events in MSNs from WT and R6/2 mice in the three age groups. At V_h_=-70 mV in ACSF, no statistically significant differences in amplitude, area, or half-amplitude duration were observed **(Suppl. Table III)**. The rise time of synaptic events was generally slower in R6/2 compared to WT MSNs. However, this difference was less apparent for glutamatergic events isolated with BIC. For GABAergic synaptic currents, we observed a significant increase in amplitude at P21 in R6/2 compared to WT MSNs (p<0.001, Mann-Whitney test). In addition, an interesting age-dependent effect was found. The amplitude of GABAergic events in WT MSNs decreased significantly from P7 to P14 (p=0.005, One-way ANOVA followed by Bonferroni post-test), and remained decreased at P21 (p=0.002). In contrast, the reduction in amplitude in R6/2 MSNs was not statistically significant between P7 and P14 and even increased back to P7 levels, suggesting a compensatory mechanism.

*Other synaptic events in developing MSNs:* In contrast to CPNs and in spite of containing all the elements necessary to generate GDPs, these synaptic events were not observed in MSNs at P7. Interestingly, a few spontaneous NMDA spikes also were recorded in MSNs **(Fig. 6B)**. They shared similar characteristics to those recorded in CPNs. With regard to paroxysmal activity, the striatum is not an epileptogenic region *per se* [however, see (Aupy et al., 2018)]. Thus, in the presence of GABA_A_ receptor blockers, epileptiform activity is rarely observed. However, other signs of hyperexcitability, e.g., large-amplitude events or bursts of spontaneous EPSCs, can become evident and they may reflect the propagation of epileptic activity from CPNs. After addition of BIC, a number of MSNs from R6/2 mice displayed paroxysmal-like events and bursts that appeared more intense than those in WT MSNs **(Suppl. Fig. 6)**, e.g., longer duration and bursting activity. In terms of MSNs morphology, biocytin labeling of MSNs was not as successful as that for CPNs, probably due to their small size. Thus, only a few examples of MSNs at the different developmental ages are illustrated **(Fig. 9)**. Dendritic abnormalities similar to those observed in CPNs were not apparent.

**Fig. 9:**
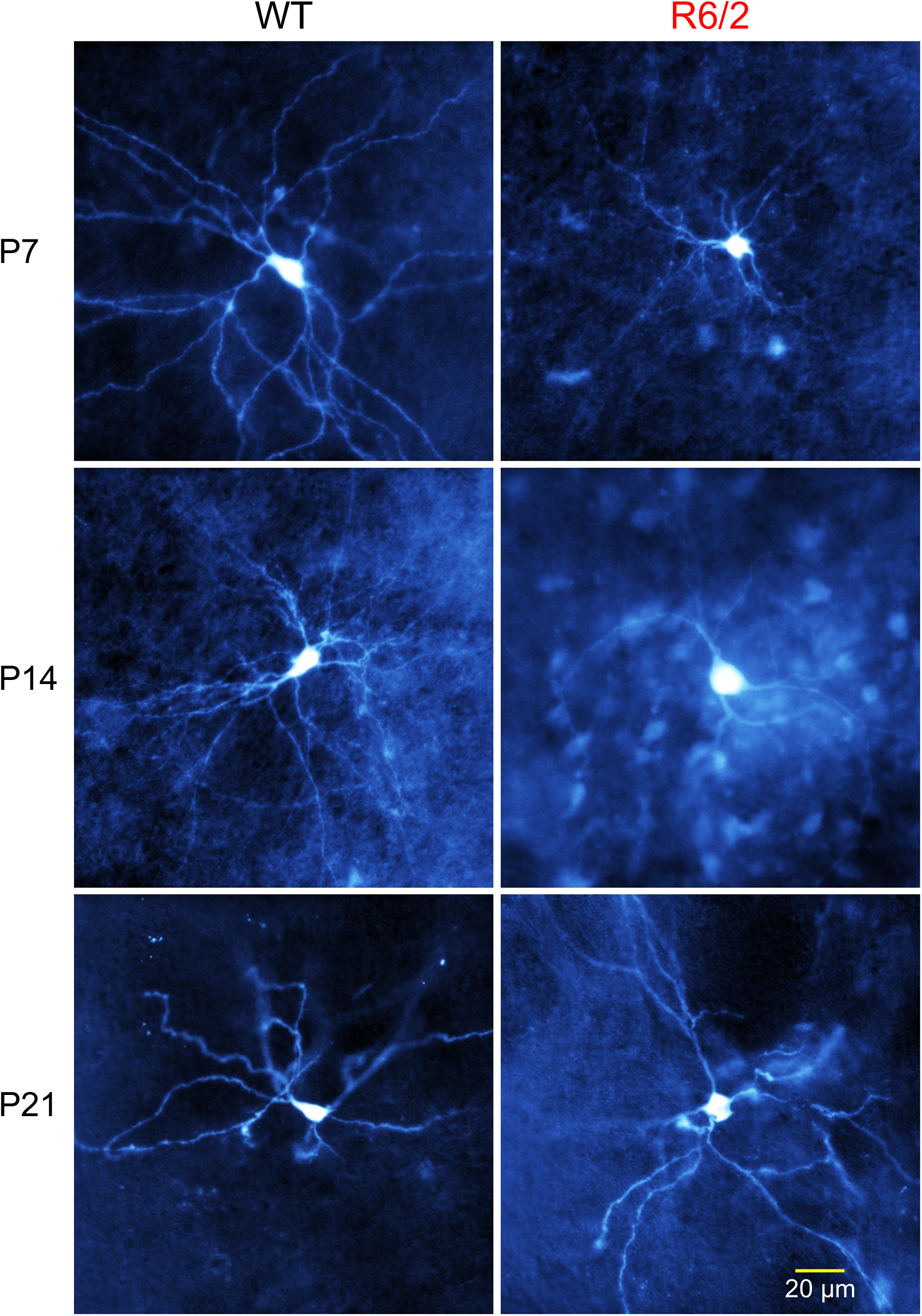
**Morphology of MSNs from WT and R6/2 mice at P7, P14, and P21**. MSNs were recorded and filled with biocytin. No major morphological differences were noticed between WT and R6/2 MSNs at any age. However, notice that at P14, the soma of the R6/2 MSN appears larger than the WT MSNs. Calibration in the bottom right applies to all panels.

## Discussion

A pressing question in HD research is to know whether HD abnormalities occur simultaneously in different regions of the brain or a specific region, e.g., striatum or cortex, triggers a cascade of events leading to circuit and system derangement. Studies have shown that cell-cell interactions play a pivotal role in HD neuropathology (Gu et al., 2005; Wang et al., 2014). However, a functional demonstration of how one region affects a connected target is still lacking. By studying the developmental trajectories of cerebral cortex and striatum in parallel, we identified cortical alterations that may contribute to striatal circuit dysfunction in R6/2 model mice. Very early during cortical development, anatomical and electrophysiological evidence demonstrates altered positioning and morphology of CPNs, general reductions in cell membrane capacitance, increases in input resistance, deficits in glutamatergic and GABAergic synaptic inputs, and enhanced presence of GDPs, all indicating delayed cortical development, at least until the second postnatal week. At 3 weeks of age, some of the alterations in intrinsic membrane and synaptic properties recover close to those of control mice. Striatal alterations in R6/2 mice are fewer, occur later than those in cortex, and often are of opposite sign, e.g., increased cell capacitance at P14. However, this does not rule out cell autonomous changes in MSNs, considering that the mHTT is ubiquitous across brain regions and in every type of cell. In fact, histopathological studies in human HD carriers appear to indicate that HTT expression affects caudate neuron densities from early in life (Myers et al., 1991). Significant alterations also occur in genetic mouse models of HD (Lebouc et al., 2020). Although some of these alterations could be due to cell-autonomous processes, there is no doubt that cortical inputs, or lack thereof, could exacerbate striatal dysfunction and eventual demise.

### Abnormal cortical architecture and morphological alterations of developing CPNs

Based on anatomical observations (i.e., NeuN staining), we demonstrate very early abnormalities of cortical development, manifested by laminar disorganization, neuronal crowding, and areas devoid of the neuronal marker, suggesting migration and neuronal deficits. Subtle at the beginning (around P14), these abnormalities become more apparent in later stages. These manifestations are reminiscent of FCD type 1 (Blumcke et al., 2011; Cepeda et al., 2006; Cepeda et al., 2019; Hickman et al., 2021; Zhang et al., 2020) and are in agreement with elegant studies showing the important role of HTT in neuronal migration and specification (Barnat et al., 2020; Godin et al., 2010; Hickman et al., 2021). More recently, using HD patient-derived human cortical organoids, altered corticogenesis was clearly demonstrated (Liu et al., 2024). It was manifested by, among others, delayed postmitotic neuronal maturation, dysregulated cortical neuron fate specification, and abnormalities in early subcortical projections, thus revealing a causal link between impaired progenitor cells and chaotic cortical lamination in HD brains. While cortical disorganization in HD brains has been demonstrated, the significance of regions devoid of the neuronal marker are more difficult to interpret. It could be argued that reduced expression of NeuN signals an ongoing degenerative processes and not neuronal disorganization or loss. In fact, using a different cell stain, e.g., cresyl violet, we were not able to demonstrate areas devoid of cells in our material, consistent with studies showing that neuronal loss in not evident even in symptomatic R6/2 mice and occurs only at very late stages of the disease (Davies et al., 1997; Turmaine et al., 2000). In agreement, it has been shown that lack of NeuN expression does not necessarily reflect neuronal vacuum (Gusel’nikova and Korzhevskiy, 2015). The most parsimonious explanation of FCD-like lesions is thus a combination of neurodevelopmental and neurodegenerative alterations (Mangin et al., 2020). This agrees with recent studies in post-mortem HD brains suggesting that cortical cell loss emerges as a consequence of neurodevelopmental changes at the beginning of life, followed by neurodegeneration in adulthood (Estevez-Fraga et al., 2023).

At the cellular level, some CPNs in R6/2 mice from P7 onward displayed overly tortuous processes and misorientation, consistent with morphological changes in adult mice from other HD models (Laforet et al., 2001) and in human patients (DiFiglia et al., 1997; Graveland et al., 1985; Sapp et al., 1999). According to the human HD studies, dystrophic neurites are more prevalent in deep cortical layers and because they could be seen in a presymptomatic adult patient, it was suggested that they could precede clinical onset (DiFiglia et al., 1997). It also has been suggested that the presence of recurved dendrites in cortical and striatal neurons may offer a clue into the pathophysiology of HD (Graveland et al., 1985). Although it is possible that these changes in adult patients are an indication of degenerative changes, we believe that dendritic and axonal abnormalities observed in developing and adult R6/2 mice, as well as in HD patients, could be a remnant of faulty development, e.g., defective pathfinding. For example, it has been shown that axonal growth is compromised in HD models due to downregulation of NUMA1 (Capizzi et al., 2022). We also observed CPN misorientation and one case of dual apical dendrites. CPN misorientation is a common finding in human FCD (Cepeda et al., 2003b) and CPNs with dual apical dendrites also have been observed in FCD (Cepeda et al., 1993) and in medial prefrontal cortex of an animal model (Wang et al., 2006). The presence of these CPNs could increase cortical excitability (Cepeda et al., 2019).

### Altered passive and active membrane properties of CPNs and MSNs

In general, the cell membrane capacitance of R6/2 CPNs at all ages was reduced and the input resistance was increased compared to WT CPNs. Reduced cell membrane capacitance became statistically significant at P14 and was still slightly reduced at P21. At P21, reductions in cell capacitance correlated well with decreased somatic area based on biocytin measurements. As expected, reduced cell capacitance was accompanied by increased input resistance (P7 and P14), likely as a result of less membrane area and functional channels. This suggests delayed CPN development and is consistent with other studies in HD mouse models (Braz et al., 2022). However, the RMP did not differ between genotypes at any age, except for a trend for more depolarized RMPs at P14 in R6/2 CPNs. In striatum, genotype differences in WT vs. R6/2 MSNs were at odds with those in CPNs. At P7, membrane capacitance of MSNs was practically identical between genotypes. Surprisingly, at P14, MSN capacitance was significantly increased in R6/2 compared to WT mice. This effect was transitory, as it did not persist at P21. Notably, we also found that the rheobase of CPNs was significantly reduced in R6/2 CPNs at P14, and this resulted in increased excitability. A transient increase in CPN excitability also was observed in Braz et al. study at P4-6 (Braz et al., 2022). CPN activity can lead to increased BDNF release from corticostriatal terminals, in particular from the motor cortex (Altar et al., 1997; Ehinger et al., 2023), which might explain increased MSN capacitance at P14. Less likely is an increase in cell capacitance due to more gap junctional communication.

### Altered spontaneous synaptic activity in CPNs and MSNs

At P7 and P14, in regular ACSF, the frequency of spontaneous synaptic currents was reduced in R6/2 CPNs. This deficit was overcome at P21. After isolation of glutamatergic inputs with BIC, the difference persisted at P7 but not in older mice, suggesting that HD brains are capable of compensating early synaptic deficits. We also showed that GDPs are more prevalent in R6/2 CPNs at P7, indicating that the GABA developmental switch (from depolarizing to hyperpolarizing), which normally occurs between the first and second postnatal week, takes longer to develop in R6/2 mice. GABA is not just a classical inhibitory neurotransmitter, during brain development GABAergic synapses develop before glutamatergic synapses (Hennou et al., 2002) and in immature networks GABA is capable of depolarizing cell membranes, activating Ca^2+^ channels, and producing a net excitatory effect (Ben-Ari, 2014; Ben-Ari and Spitzer, 2004) due to high expression of NKCC1 chloride transporter and reduced expression of KCC2 (Rivera et al., 1999; Sedmak et al., 2016; Virtanen et al., 2020). GABA also plays a critical role as a trophic factor, supporting cell migration and differentiation (Kriegstein and Owens, 2001; Owens and Kriegstein, 2002; Represa and Ben-Ari, 2005). If, as we showed, the GABA developmental switch is abnormal in HD brains, this also could contribute to the generation of FCD-like areas in the cortex.

The striatal circuitry develops later than the cortical circuit and is highly dependent on cortical inputs. For a review on striatal development in normal and HD conditions see (Lebouc et al., 2020; Tepper et al., 1998). Interestingly, in spite of delayed maturation of cortical inputs in R6/2 mice, no deficits in MSN spontaneous glutamatergic activity were observed at P7 and P14. Although CPNs start sending their axonal projections to the striatum from P3 (Sohur et al., 2014), in a previous study we showed that only a few MSNs respond to cortical stimulation during the first postnatal week and, when responses are evoked, most of the current is mediated by non-NMDA receptors (Colwell et al., 1998; Hurst et al., 2001). During the second postnatal week, a strengthening of corticostriatal synapses takes place and the contribution of NMDA receptors increases (Colwell et al., 1998; Hurst et al., 2001; Krajeski et al., 2019). However, thalamostriatal projections seem to develop earlier, as VGLUT2-positive axons are present before VGLUT1- positive axons in the mouse telencephalic and diencephalic regions and are already found at birth in the mouse striatum (Nakamura et al., 2005). In consequence, it is likely that the absence of glutamatergic deficits in P7 MSNs is the result of a more mature thalamic input. Alterations in thalamocortical, and also presumably thalamostriatal, projections occur late in HD mice (Holley et al., 2022). Although previous anatomical studies in the Q140 knock-in model of HD demonstrated that the loss of thalamostriatal terminals precedes the loss of corticostriatal projections (Deng et al., 2014), the early development or loss of thalamostriatal synapses in R6/2 mice has not yet been examined, in particular the Cm/Pf projection. Our studies demonstrated altered thalamocortical communication in symptomatic but not in pre-symptomatic R6/2 mice. Although some changes in intrinsic properties in VAL and VPM thalamic nuclei were observed at P21, the input-output functions were not significantly altered, suggesting that the thalamic output, at least from these thalamic nuclei, is not affected (Holley et al., 2022). However, by P21 glutamatergic inputs start to decrease in R6/2 mice and this deficit becomes more pronounced in later stages of disease progression, when motor symptoms appear (around 5-7 weeks) (Cepeda et al., 2003a). Thus, it appears that the changes in excitatory inputs in CPNs and MSNs are divergent. While reduced glutamatergic inputs in CPNs eventually recover, those in MSNs, which were normal at P7, progressively deteriorate.

An important issue remains unexplained; why does the transient increase in CPN excitability at P14 not translate into increased glutamatergic inputs to MSNs? One possibility is that MSNs are able to engage homeostatic mechanisms to reduce corticostriatal drive, which could be potentially harmful to the cell (Turrigiano, 2012). These may include presynaptic modulation of glutamate release *via* dopamine D2 receptors (Bamford et al., 2004; Cepeda et al., 2001b), GABA_B_ receptors (Calabresi et al., 1990; Logie et al., 2013), or endocannabinoid receptors (Gerdeman and Lovinger, 2001) known to be present in corticostriatal terminals.

Interestingly, in striatal MSNs there was a significant deficit of GABAergic inputs at P14. As the main GABAergic input onto MSNs is from local interneurons, it is possible that PV- and SOM-expressing interneurons also are immature. Additional studies on the development of GABAergic interneurons in cortex and striatum should be performed in order to answer this question. However, by P21 this GABAergic deficit is also reversed by P21 and with disease progression their frequency exceeds that of WT MSNs (Cepeda et al., 2004). Thus, P14 appears to be a critical age or inflexion point in terms of the future fate of excitatory and inhibitory synaptic dysfunction in both cortex and striatum. For example, as shown here, a deficiency of glutamatergic synaptic inputs to MSNs first occurs during this age range (P14-P21). This may be due to reduced cortical and/or thalamic inputs, increased synaptic pruning, and decreased glutamate release probability from existing terminals. Remarkably, in a cell model in which YAC128 (a full-length HD model) and WT striatal neurons were co-cultured with cortical neurons, it was demonstrated that corticostriatal connectivity in WTs develops rapidly and continuously from 7 to 21 days *in vitro* (DIV), whereas YAC128 connectivity showed no significant growth from 14 DIV onwards. This effect implicated a reduction in post-synaptic dendritic arborization and the size and replenishment rate of the presynaptic readily releasable pool of excitatory vesicles (Buren et al., 2016). Further, in a model of autism spectrum disorders, accelerated maturation and remodeling of corticostriatal circuits during the second and third postnatal weeks occurred as a result of cortical hyperactivity (Peixoto et al., 2016).

The role of synaptic and extrasynaptic NMDA receptors in HD is well established (Cepeda et al., 2001a; Levine et al., 1999; Milnerwood et al., 2010; Raymond et al., 2011; Zeron et al., 2002). How NMDA receptors develop in WT vs. HD mice is of great importance. Although in the present study we could not separate glutamatergic activity by receptor subtype, we found that a small percentage of CPNs and MSNs (∼12-20% regardless of age or genotype) displayed rare, spontaneous NMDA spikes at P14 and P21 in ACSF and after the addition of BIC. Although their amplitude was equivalent between genotypes, their duration was slightly increased in CPNs from R6/2 mice. NMDA spikes are regenerative potentials generated locally in the basal dendrites of CPNs and serve to amplify synaptic input (Antic et al., 2010; Chalifoux and Carter, 2011; Kumar et al., 2018; Schiller et al., 2000). Their significance in diseased states is just beginning to be explored. In adult YAC128 mice, NMDA spikes could be evoked in CPNs and more neurons displayed these large amplitude NMDA events (Sepers et al., 2022). Large amplitude glutamatergic events in R6/2 mice were first described by our group and, in retrospect, some of these were in fact NMDA spikes [see Fig. 1 in (Cepeda et al., 2003a)]. The occurrence of spontaneous NMDA spikes has been reported anecdotally (Kumar et al., 2018; Oikonomou et al., 2015; Sepers et al., 2022), but a clear demonstration of their existence has been elusive, particularly in MSNs, although models of striatal synaptic integration predicted their existence (Lindroos and Hellgren Kotaleski, 2021). It also was shown recently that NMDA spikes in MSNs can be facilitated by the depolarizing actions of GABA (Day et al., 2024). Considering that in R6/2 MSNs GABA synaptic activity progressively increases as mice approach the overt phenotype (Cepeda et al., 2004), we could speculate that the interplay between depolarizing actions of GABA, increased membrane input resistance (due to dendritic damage and spine loss), and more pervasive localization of NMDA receptors at extrasynaptic sites, facilitates the generation of NMDA spikes, leading to further deleterious effects.

### Consequences of increased cortical excitability

In a previous paper we hypothesized that cortical maldevelopment is the *primum movens* of excitatory/inhibitory imbalances in HD (Cepeda et al., 2019). With HD progression, CPNs become hyperexcitable probably due to a combination of FCD-like alterations (aberrant lamination, misorientation, tortuous dendrites and axons), neurodegenerative changes (loss of spines and overall membrane area), and dysfunction or loss of PV-expressing interneurons (Burgold et al., 2019; Kim et al., 2014). GABA synaptic transmission in R6/2 mice is deficient during early cortical development (P7 and to some extent P14) but, with age, it eventually becomes similar or even upregulated compared to that in control mice (Cummings et al., 2009). Further, in a late-onset knock-in model of HD (Q175), an increase in GABA synaptic transmission in the cortex was observed before full manifestation of symptoms, probably to counter increased cortical excitability (Indersmitten et al., 2015). In developing R6/2 mice spontaneous seizures do not occur, however, blockade of GABA_A_ receptors unveils cortical hyperexcitability, manifested by longer duration and more complex morphology of PDs [(see also (Cummings et al., 2009)]. Nevertheless, in spite of accumulating evidence that brain development in HD patients and animal models is abnormal and the cortex becomes hyperexcitable (Cepeda et al., 2019), the phenotype does not manifest immediately. In the case of human adult-onset HD, it takes many years before overt symptoms become evident. One possible explanation, as stated above, is that the HD brain is capable of compensation to cope with early developmental abnormalities [see (Mackay et al., 2018; Oikonomou et al., 2021)]. However, when cortical inhibition falters, symptoms emerge, starting with cognitive and psychiatric disturbances followed by motor symptoms. Thus, the onset of the HD phenotype could be the consequence of failure of cortical inhibition in a hyperexcitable cortex. In contrast, early deficits in striatal GABA transmission, evolve into an upregulation of inhibitory activity (Cepeda et al., 2004).

### Insights into HD therapies

Based on recent studies in human HD and in animal models, there is no doubt that neurodevelopment is altered, implying that any therapeutic intervention attempting to reverse or ameliorate HD symptoms has to start very early (Cepeda et al., 2019). Indeed, in a knock-in model of adult-onset HD (Hdh^Q111/Q7^), systemic administration of an ampakine (CX516) during the first postnatal week increases AMPA receptor function, restores dendritic arborization complexity, and improves sensorimotor function in HD pups (Braz et al., 2022). This study also showed that even without intervention, HD pups are capable of spontaneous rescue of synaptic deficits, even though they later develop the phenotype. The question is, should we try to avoid self-restoring mechanisms and intervene before these intrinsic compensatory processes take place? The fact that HD brains are capable of autocorrection poses an interesting dilemma. Considering that both mouse and humans are altricial animals and that the first postnatal week in mice corresponds roughly to the third trimester of embryonic life (Clancy et al., 2007; Zeiss, 2021), this type of treatment would have to be applied *in utero*. Another consideration is that, depending on the dose, ampakines in R6/2 mice may increase cortical excitability beyond normal limits and exacerbate a pro-epileptic state (Cepeda et al., 2010). In fact, in the Braz et al. study, CX516 had deleterious effects in WT mice (Braz et al., 2022). It also is important to notice that the motor phenotype in this model starts around 9 months of age (Yhnell et al., 2016), corticostriatal synaptic deficits are only observed around 4-5 months, and subtle brain volume and cortical thinning can be found at 10 months but are less apparent at 18 months of age (Kovalenko et al., 2018). As ampakines also increase BDNF (Simmons et al., 2009), beneficial effects in mouse pups could be better explained by their pleiotropic effects on neuronal structure and synaptic activity. For example, we demonstrated that BDNF can reverse increases in GABA synaptic activity of symptomatic R6/2 MSNs (Cepeda et al., 2004). In addition, BDNF can also regulate chloride transport through the modulation of the chloride transporter KCC2 (Hamze et al., 2024).

In a study using HD-derived induced pluripotent stem cell lines, it was shown that early telencephalic induction and late neural identity are affected in cortical and striatal populations (Conforti et al., 2018). Thus, large CAG expansions led to failure of neuro-ectodermal acquisition, while cells carrying shorter CAGs repeats showed gross abnormalities in neural rosette formation as well as disrupted cytoarchitecture in cortical organoids. Importantly, HD organoids displayed an immature transcriptional blueprint as demonstrated by downregulation of transcripts involved in neuronal migration and differentiation. Further, these transcripts also were downregulated in embryonic cortex (E16.5) from R6/2 model mice, supporting incomplete or delayed maturation. Notably, these defects could be rescued by downregulation of mHTT or pharmacological inhibition of ADAM10. In another model of HD using human cortical organoids from control (hCOs) and HD patients (HD-hCOs), alterations of corticogenesis in HD-hCOs were found (Liu et al., 2024). Pathology included deficient junctional complexes, premature neuronal differentiation, and delayed neuronal maturation, revealing a causal link between impaired progenitor cells and chaotic cortical neuronal layering in the HD brains. Remarkably, the authors suggested that restoring the recruitment of ADP-ribosylation factor 1 (ARF1) to Golgi in HD neural tubes might rescue the altered corticogenesis in HD fetal brain (Liu et al., 2024).

## Conclusions

Cortical development is a complex and delicate process that follows precise developmental pathways or chreods (Waddington, 1962). For that reason, it is easily exposed to external and internal perturbations. The present contribution adds to the already extensive literature on developmental alterations due to the presence of mHTT. Very early on, morphological and functional deviations from normal development occur in the cerebral cortex of R6/2 mice. Some of these changes are autocorrected, while others are more permanent. In fact, HD brains appear to undergo homeorhesis, in an attempt to return to its normal developmental trajectory. Discovering and promoting intrinsic compensatory mechanisms, in conjunction with early intervention, will aid in finding the best therapeutic approaches for HD.

## CRediT authorship contribution statement

**Carlos Cepeda:** Writing – review & editing, Writing – original draft, Visualization, Validation, Supervision, Investigation, Formal analysis, Conceptualization. **Sandra M. Holley:** Writing – review & editing, Investigation, Formal analysis. **Joshua Barry:** Writing – review & editing, Investigation, Formal analysis. **Katerina D. Oikonomou:** Writing – review & editing, Investigation, Formal analysis. **Vannah-Wila Yazon:** Writing – review & editing, Investigation, Formal analysis. **Allison Peng:** Writing – review & editing, Formal analysis, Investigation. **Deneen Argueta:** Writing – review & editing, Formal analysis, **Michael S. Levine:** Writing – review & editing, Visualization, Supervision, Conceptualization.

## Data Availability

Original data can be provided by the corresponding author upon reasonable request.

## Acknowledgments

We would like to acknowledge the participation of Elissa Donzis, Samiksha Chopra, Minh Bui, Jan Asai, in data acquisition and analyses. We also appreciate insightful comments from Dr. Joseph Watson. This work was supported by USPHS grant NS111316 (CC).

## Supplementary Figure

**Suppl. Fig. 1.**
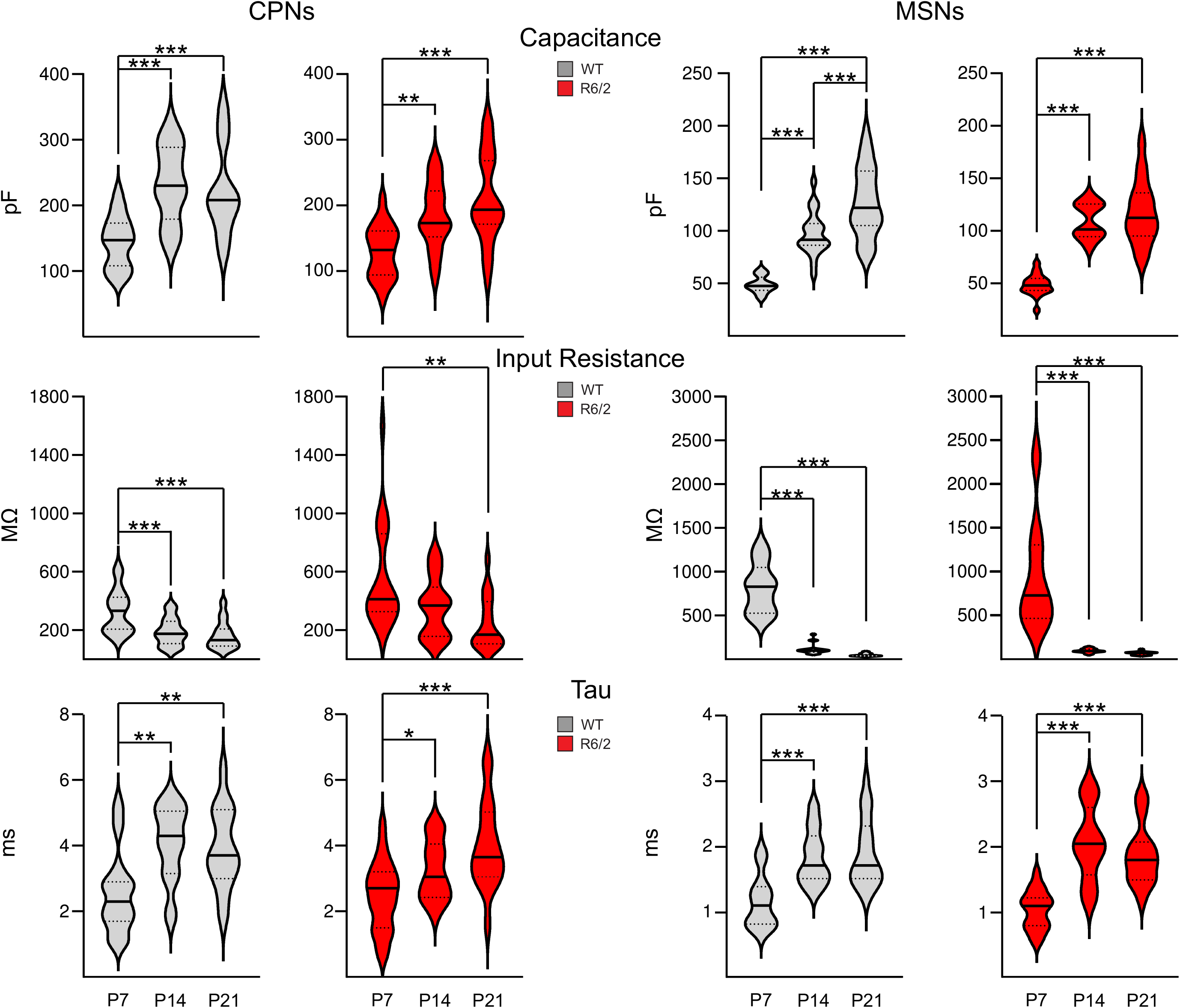
: Developmental trajectory of passive membrane properties of CPNs and MSNs. Cell membrane properties (capacitance, input resistance and decay time constant) of CPNs and MSNs were measured in voltage clamp mode with Cs-methanesulfonate as the internal pipette solution in WT and R6/2 mice at P7, P14, and P21. Violin plots show the median (solid line), lower and upper quartiles (dotted lines), as well as the distributions of numerical data points using density curves. Data were analyzed using One-way ANOVA followed by multiple comparisons (Bonferroni *post hoc* test). In this and subsequent figures, asterisks indicate statistically significant differences; * p<0.05, ** p<0.01, and *** p<0.001.

**Suppl. Fig. 2.**
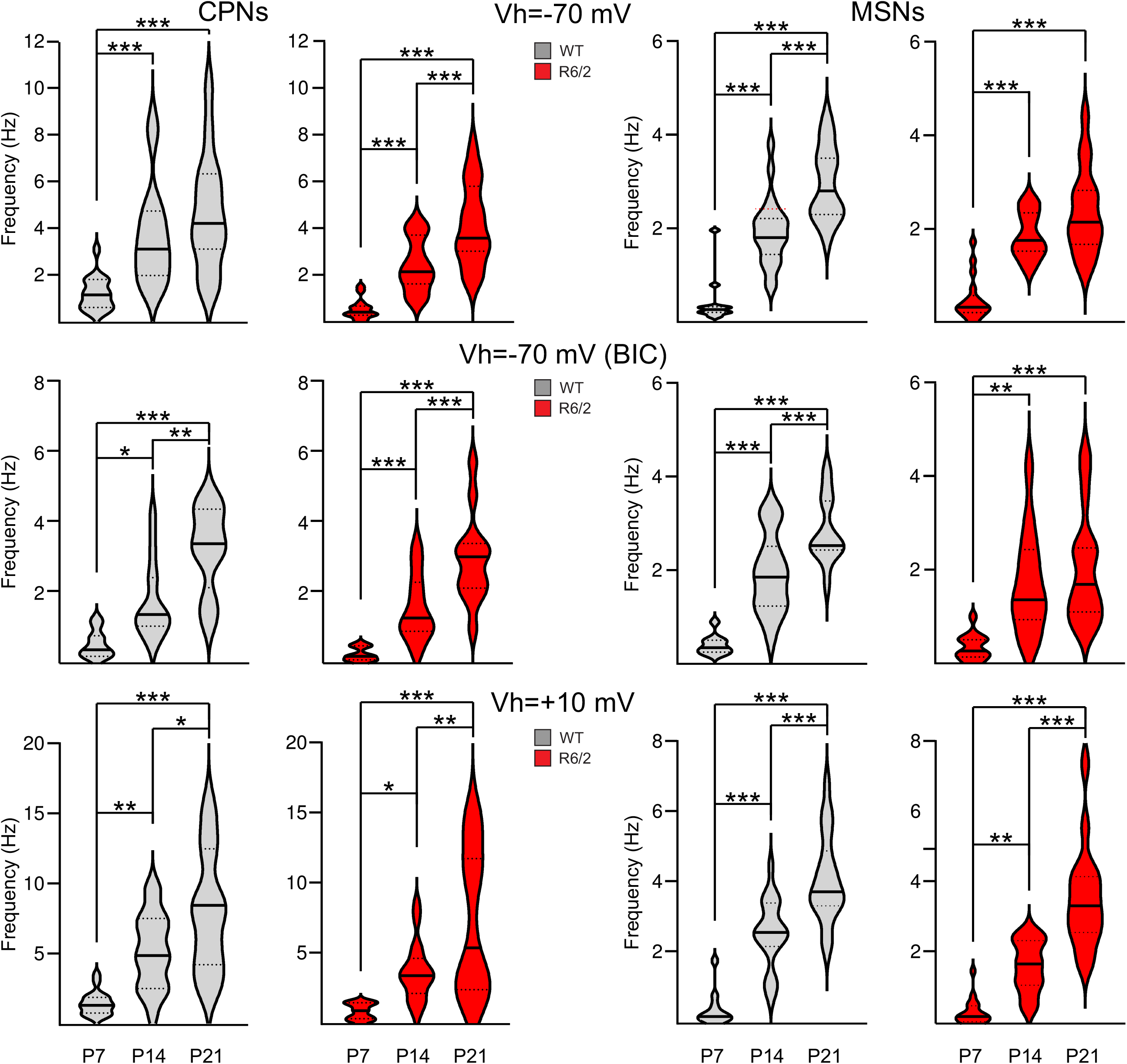
: Developmental trajectory of spontaneous synaptic activity of CPNs and MSNs. Synaptic currents were recorded in voltage clamp mode with Cs-methanesulfonate as the internal pipette solution in WT and R6/2 mice at P7, P14, and P21. Violin plots show the median (solid line), lower and upper quartiles (dotted lines), as well as the distributions of numerical data points using density curves. Data were analyzed using One-way ANOVA followed by multiple comparisons (Bonferroni *post hoc* test).

**Suppl. Fig. 3.**
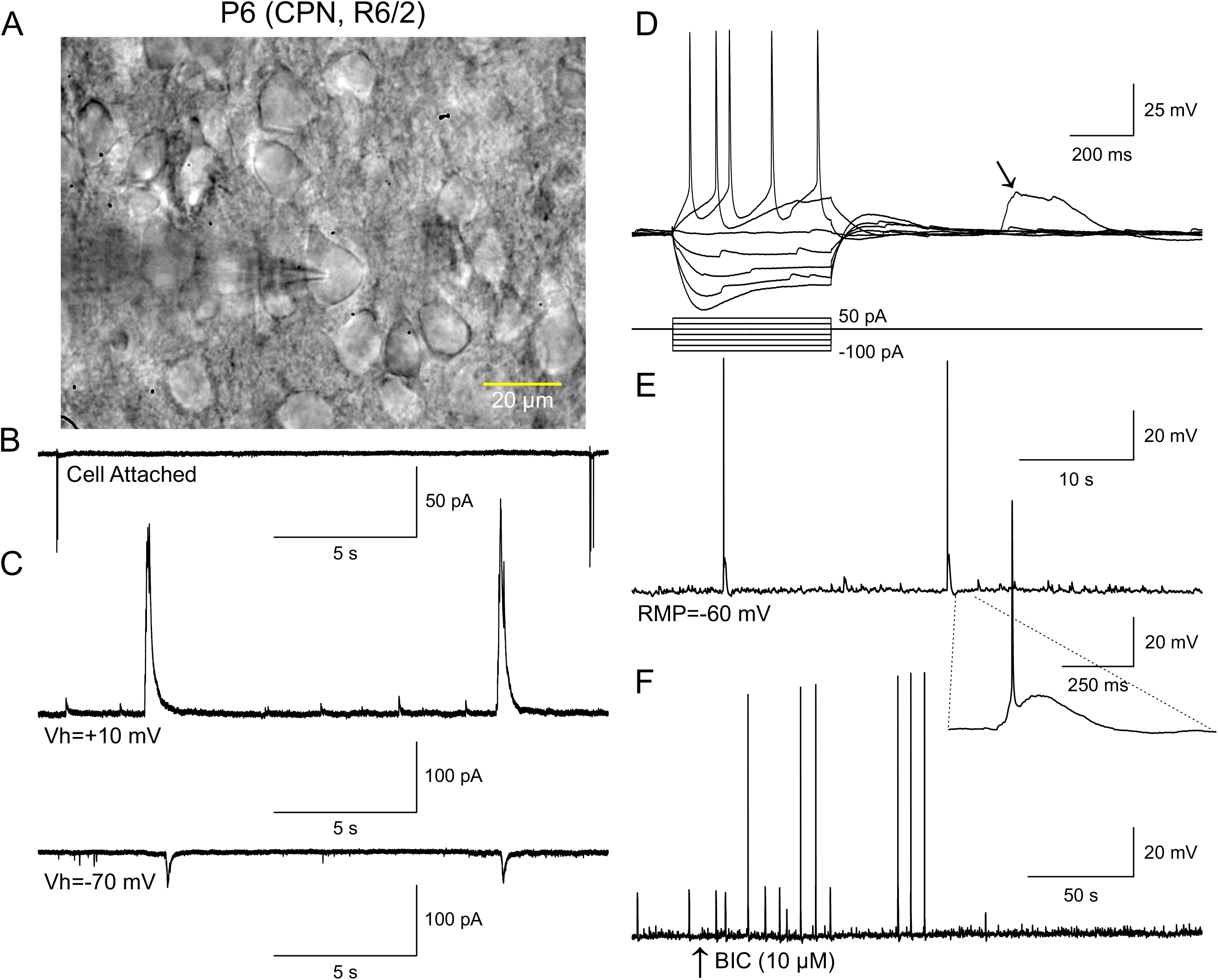
: Spontaneous GDPs recorded in CPNs of R6/2 mice at P7. **(A)** IR-DIC image of a CPN with the patch electrode attached. **(B)** In this cell, rhythmic, spontaneous action potentials were observed before opening the gigaohm seal (cell attached mode). This indicates that the GDPs depolarized the cell membrane to the threshold for firing. **(C)** After opening the seal, spontaneous GABA currents were recorded at V_h_=-70 mV and V_h_=+10 mV. **(D)** In another CPN, spontaneous GDPs were observed in current clamp mode (K-gluconate in the internal patch pipette). This GDP (arrow) occurred after hyperpolarizing and depolarizing current pulses were injected in the cell, which showed typical responses of CPNs. **(E)** Some GDPs in the same cell reached the threshold for firing action potentials. **(F)** After addition of BIC (10 µM), GDPs were abolished after 1-2 min application.

**Suppl. Fig. 4.**
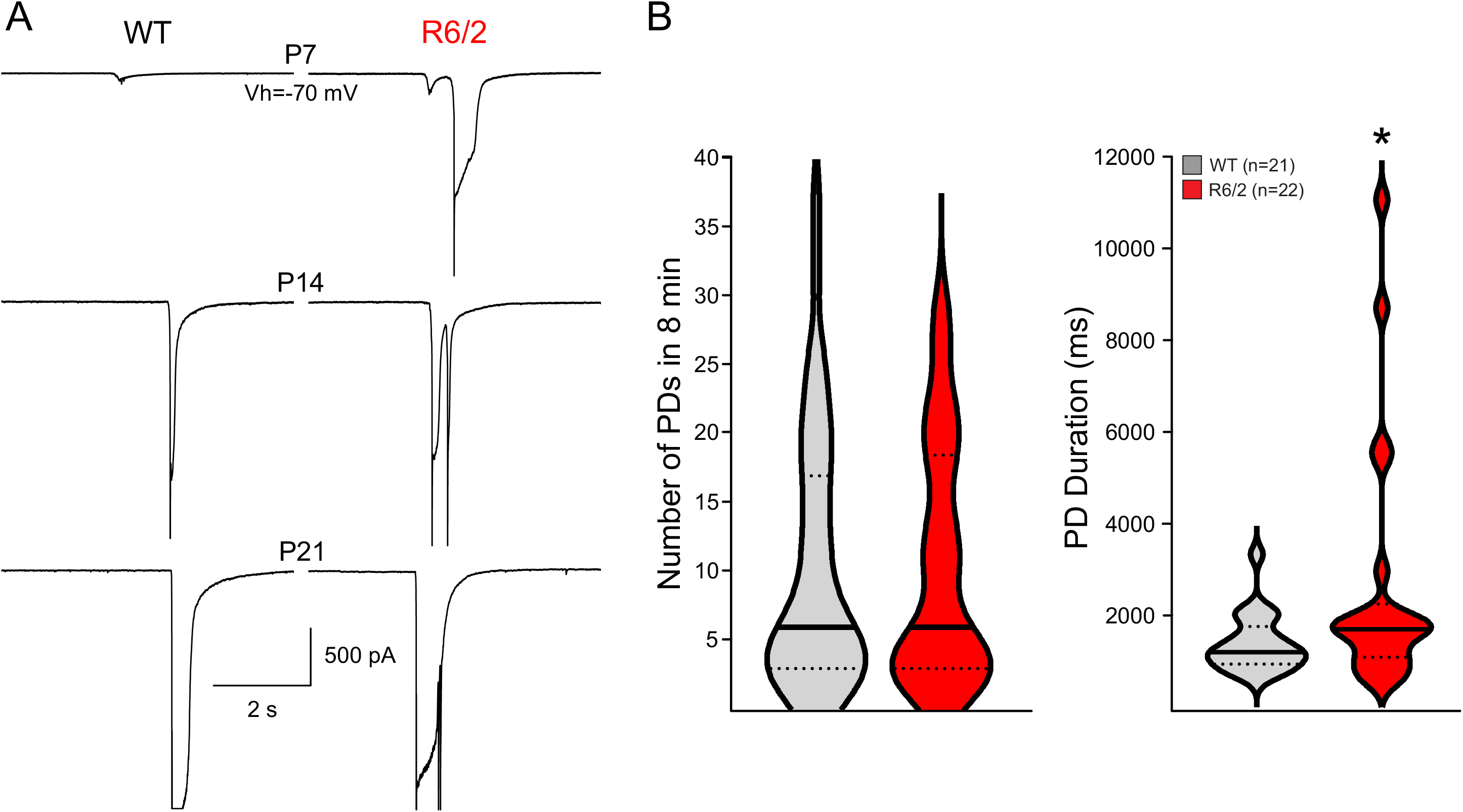
: The duration of PDs in CPNs is increased in R6/2 mice. Paroxysmal activity was induced in CPNs by adding BIC (10 µM) to the bath solution. In the graphs, PDs recorded in WT and R6/2 CPNs at P7, P14, and P21 were combined. **(A)** Representative traces of PDs evoked in WT and R6/2 CPNs at P7, P14, and P21 (V_h_=-70 mV). Notice that PDs in R6/2 CPNs were more complex and had increased duration compared to WT CPNs. **(B)** Violin plots show the median (solid lines) frequency and duration of PDs in WT and R6/2 CPNs. While there was no difference in the number of PDs, their duration was increased in R6/2 CPNs (p=0.049, Student’s t-test).

**Suppl. Fig. 5.**
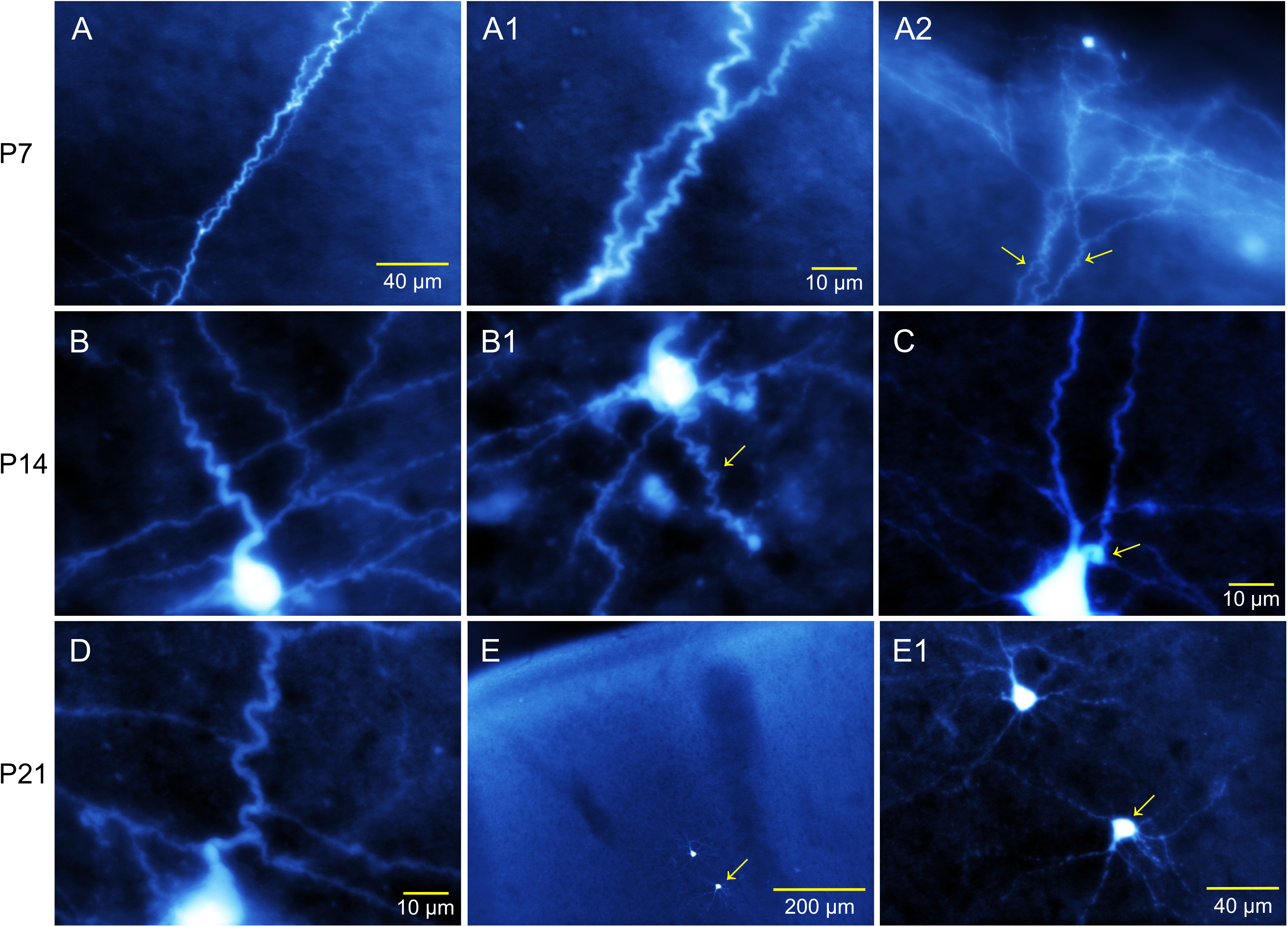
: Morphological cellular abnormalities of CPNs in R6/2 mice. High magnification images (A-D) of biocytin-filled CPNs. **(A, A1, A2).** This CPN recorded at P7 showed excessively meandering apical dendrite **(A, A1)** and terminal tuft. **(B, B1)** CPN recorded at P14 (also shown in Fig. 7, right panel) also displaying overly tortuous processes, including a sharp bend close to the soma, tortuous apical dendrite **(B)** and axon **(B1)**. **(C)** CPN at P14 showing what appears to be two apical dendrites emerging from the soma. The second dendrite also has a sharp bend (arrow). **(D)** Another CPN at P21 displaying a sharp bend close to the soma and tortuous apical dendrite. **(E, E1)** CPNs demonstrating the existence of misorientation in R6/2 cortex. The low magnification image **(E)** shows two CPNs (P21), the higher magnification image **(E1)** shows that the bottom cell is oriented opposite to the cortical surface.

**Suppl. Fig. 6.**
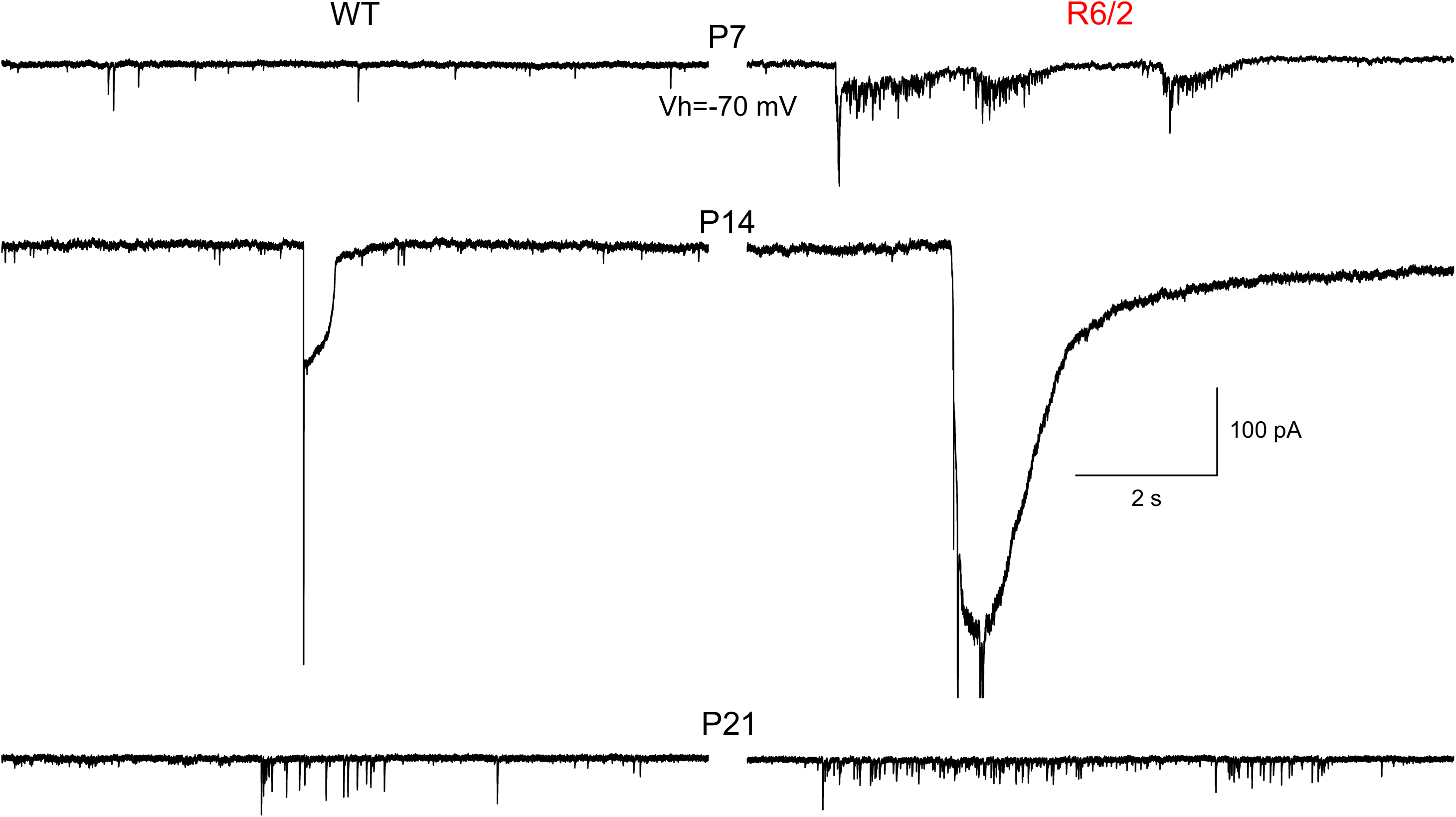
: **Propagation of epileptiform activity to MSNs in WT and R6/2 mice.** MSNs were recorded in the presence of BIC (10 µM) at P7, P14, and P21. Bath application of BIC induced paroxysmal discharges in CPNs (not shown, as both regions were not examined simultaneously). This activity occasionally propagated to the striatal MSNs. Cells from R6/2 mice displayed more excitability than WT MSNs, manifested by bursts of glutamatergic activity and in some cases by large inward, paroxysmal-like discharges (e.g., at P14).

